# Autosomal Recessive Alzheimer’s disease (arAD): homozygosity mapping of genomic regions containing arAD loci

**DOI:** 10.1101/2020.02.10.941393

**Authors:** Sonia Moreno-Grau, Maria Victoria Fernández, Itziar de Rojas, Isabel Hernández, Fabiana Farias, John P Budde, Inés Quintela, Laura Madrid, Antonio González-Perez, Laura Montrreal, Pablo Garcia-Gonzalez, Emilio Alarcón-Martín, Montserrat Alegret, Olalla Maroñas, Juan Antonio Pineda, Juan Macías, GR@ACE & DEGESCO consortia, Alzheimer’s Disease Neuroimaging Initiative, Marta Marquié, Sergi Valero, Alba Benaque, Jordi Clarimón, Maria Jesus Bullido, Guillermo García-Ribas, Pau Pástor, Pascual Sánchez-Juan, Victoria Álvarez, Gerard Piñol-Ripoll, Jose María García-Alberca, José Luis Royo, Emilio Franco-Macías, Pablo Mir, Miguel Calero, Miguel Medina, Alberto Rábano, Jesús Ávila, Carmen Antúnez, Luis Miguel Real, Adelina Orellana, Ángel Carracedo, María Eugenia Sáez, Lluis Tárraga, Mercè Boada, Carlos Cruchaga, Agustín Ruiz

**Affiliations:** Research Center and Memory clinic Fundació ACE. Institut Català de Neurociències Aplicades. Universitat Internacional de Catalunya, Barcelona, Spain; CIBERNED, Center for Networked Biomedical Research on Neurodegenerative Diseases, Carlos III Institute of Health, Spain; Department of Psychiatry, Washington University School of Medicine, St. Louis, MO, United States of America; Hope Center for Neurological Disorders, Washington University School of Medicine, St. Louis, MO, United States of America; Grupo de Medicina Xenómica, Centro Nacional de Genotipado (CEGEN-PRB3-ISCIII). Universidade de Santiago de Compostela, Santiago de Compostela, Spain; CAEBI. Centro Andaluz de Estudios Bioinformáticos, Sevilla, Spain; Unidad Clínica de Enfermedades Infecciosas y Microbiología. Hospital Universitario de Valme, Sevilla, Spain; Memory Unit, Neurology Department and Sant Pau Biomedical Research Institute, Hospital de la Santa Creu i Sant Pau, Universitat Autònoma de Barcelona, Barcelona, Spain; Centro de Biologia Molecular Severo Ochoa (C.S.I.C.-U.A.M.), Universidad Autonoma de Madrid, Madrid, Spain; Instituto de Investigacion Sanitaria “Hospital la Paz” (IdIPaz), Madrid, Spain; Hospital Universitario Ramón y Cajal, Madrid, Spain; Fundació per la Recerca Biomèdica i Social Mútua Terrassa, and Memory Disorders Unit, Department of Neurology, Hospital Universitari Mutua de Terrassa, University of Barcelona School of Medicine, Terrassa, Barcelona, Spain; Neurology Service “Marqués de Valdecilla” University Hospital (University of Cantabria and IDIVAL), Santander, Spain; Laboratorio de Genética Hospital Universitario Central de Asturias, Oviedo; Instituto de Investigación Biosanitaria del Principado de Asturias (ISPA); Unitat Trastorns Cognitius, Hospital Universitari Santa Maria de Lleida, Institut de Recerca Biomédica de Lleida (IRBLLeida), Lleida, España; Alzheimer Research Center & Memory Clinic, Andalusian Institute for Neuroscience, Málaga, Spain; Dep. of Surgery, Biochemistry and Molecular Biology, School of Medicine. University of Málaga, Málaga, Spain; Unidad de Demencias, Servicio de Neurología y Neurofisiología. Instituto de Biomedicina de Sevilla (IBiS), Hospital Universitario Virgen del Rocío/CSIC/Universidad de Sevilla, Seville, Spain; Unidad de Trastornos del Movimiento, Servicio de Neurología y Neurofisiología. Instituto de Biomedicina de Sevilla (IBiS), Hospital Universitario Virgen del Rocío/CSIC/Universidad de Sevilla, Seville, Spain; CIEN Foundation, Queen Sofia Foundation Alzheimer Center, Madrid, Spain; Instituto de Salud Carlos III (ISCIII), Madrid, Spain; BT-CIEN; Department of Molecular Neuropathology, Centro de Biología Molecular “Severo Ochoa” (CBMSO), Consejo Superior de Investigaciones Científicas (CSIC)/Universidad Autónoma de Madrid (UAM); Unidad de Demencias, Hospital Clínico Universitario Virgen de la Arrixaca; Fundación Pública Galega de Medicina Xenómica-CIBERER-IDIS, Santiago de Compostela, Spain

**Author notes:** Corresponding author: Agustín Ruiz M.D. Ph.D., Address: Research Center. Fundació ACE. Institut Català de Neurociències Aplicades. C/ Marquès de Sentmenat, 57, 08029 Barcelona, Spain, Email id. **Alzheimer’s Disease Neuroimaging Initiative**: Data used in preparing this article were obtained from the Alzheimer’s Disease Neuroimaging Initiative (ADNI) database (adni.loni.usc.edu). As such, the investigators within the ADNI contributed to the design and implementation of ADNI and/or provided data but did not participate in the analysis or writing of this report. A complete listing of ADNI investigators can be found at http://adni.loni.usc.edu/wp-content/uploads/how_to_apply/ADNI_Acknowledgement_List.pdf.

## Abstract

Long runs of homozygosity (ROH) are contiguous stretches of homozygous genotypes, which are a footprint of recent inbreeding and recessive inheritance. The presence of recessive loci is suggested for Alzheimer’s disease (AD). However, the search for recessive variants has been poorly assessed to date. To investigate homozygosity in AD, we performed a fine-scale ROH analysis including 21,100 individuals from 10 cohorts of European ancestry (11,919 AD cases and 9,181 controls). We detected an increase of homozygosity in AD cases compared to controls [β_FROH_ (CI95%) = 0.051 (0.023 – 0.078); P = 3.25 x 10^-4^]. ROHs increasing the risk of AD (OR > 1) were significantly overrepresented compared to ROHs increasing protection (p < 2.20 x 10^-16^). The top associated ROH with AD risk (β (CI95%) = 1.09 (0.48 ‒ 1.48), p value = 9.03 x 10^-4^) was detected upstream the *HS3ST1* locus (chr4:11,189,482‒11,305,456), previously related to AD. Next, to construct a homozygosity map of AD cases, we selected ROHs shared by inbred AD cases extracted from an outbred population. We used whole-exome sequencing data from 1,449 individuals from the Knight-ADRC-NIA-LOAD (KANL) cohort to identify potential recessive variants in candidate ROHs. We detected a candidate marker, rs117458494, mapped in the *SPON1* locus, which has been previously associated with amyloid metabolism. Here, we provide a research framework to look for recessive variants in AD using outbred populations. Our results showed that AD cases have enriched homozygosity, suggesting that recessive effects may explain a proportion of AD heritability.

## Introduction

Alzheimer’s disease (AD) is a neurodegenerative disorder that is the leading cause of dementia worldwide ^1^. A small proportion of patients develop AD before the age of 65; this is known as early-onset AD (EOAD). In most persons, clinical symptoms begin after the age of 65, in a form of the disorder known as late-onset AD (LOAD). AD presents a strong genetic component. In fact, heritability estimations for EOAD and LOAD fall in the range of 92 to 100% and 13 to 73%, respectively ^2, 3^.

Specific autosomal dominant mutations have been linked to familial EOAD: mutations in *presenilin 1* (*PSEN1*) ^4^, *presenilin 2* (*PSEN2*) ^5^, and *amyloid precursor protein* (*APP)* ^6^. These findings were pivotal events pinpointing to the role of amyloid metabolism as a disease-causing mechanism ^7^. Despite that, dominant causes account for a minority of both familial and apparently sporadic EOAD cases. It has been suggested that autosomal recessive loci might cause most EOAD cases (∼90%) ^2^. However, only two recessive mutations in the *APP* gene (A673V and E693Δ) have been described to date ^8, 9^, and these mode of inheritance remain controversial.

The most common AD clinical presentation, the sporadic form of LOAD, has a polygenic background. Genome-wide association studies (GWAS) and large sequencing projects have identified nearly 40 genetic variants associated with LOAD risk ^10, 11, 12, 13^. These discoveries only explain a limited part of disease heritability (⁓31%) ^14^. Current genetic findings were made using an additive mode of inheritance, which overlooks the relevance of non-additive genetic components, i.e. the recessive model. Despite the fact these components could explain a part of disease heritability.

It is well known that inbreeding increases the incidence of recessive diseases. Hence, the probability of detecting a recessive locus increases in offspring of consanguineous unions ^15^ because the partners share alleles inherited from a recent common ancestor. This recent parental relatedness points to genuine regions of autozygosity. Long runs of homozygosity (ROHs) — long stretches of consecutive homozygous genotypes (>1 Mb) — are a recognized signature of recessive inheritance. Thus far, they have been used for homozygosity mapping ^16^. Population history, e.g. historical bottlenecks or geographical isolation, also influences homozygosity levels in individual genomes ^17, 18^.

To assess the role of recessive inheritance in AD, Farrer et al. ^19^ studied 183 families of the isolated Wadi Ara region (an area in Israel populated mainly by Arab citizens). The Wadi Ara population has increased parental relatedness and a high prevalence of AD. Farrer et al. pointed to candidate regions with potential recessive loci ^19, 20^. Using homozygosity mapping in a consanguineous EOAD family, and subsequent sequencing of candidate regions, Bras et al. suggested the *CTFS* gene as a potential recessive locus ^21, 22^.

It has recently been demonstrated that ROHs are ubiquitous even in outbred populations ^23, 24^. An excess of homozygosity has been associated with risk of AD in individuals of Caribbean-Hispanic and African-American ancestries ^25, 26, 27^. It suggests the presence of inbreeding and potentially autosomal recessive AD (arAD) cases nested in these populations. Conversely, this association presented controversial results for individuals of European ancestry ^28, 29^. Several factors might explain these inconsistencies. First, ROH patterns differ between populations. Specific recent bottlenecks, as well as the presence of cultural practices promoting endogamous marriages in Latino groups, could be increasing inbreeding, and consequently ROH estimations, in these populations ^30, 31^. Second, it has been estimated that large sample sizes (12,000‒65,000) are required to detect an excess of homozygosity in outbred populations ^32^. Thus, previous studies might be underpowered.

Assessing the impact of inbreeding in the genetic architecture of AD remains a challenge. The limited number of deeply characterized consanguineous families, the difficulties in finding familial information for sporadic AD individuals (mainly due to the late onset of the disease) and the reduced size of intragenerational pedigrees in western countries make the search for arAD loci complex. Furthermore, follow-up of candidate ROHs in sequencing data might be a necessary step in the definitive mapping a recessive locus, but it has been poorly assessed to date. Considering limitations, we think that capturing the fraction of consanguineous individuals nested in AD cases in an outbred population could be an efficient strategy to prioritize homozygous regions potentially harboring recessive loci.

To the best of our knowledge, this is the largest genomic data set exploring the influence of homozygosity in AD (n = 21,100). First, we investigated whether AD individuals from a European outbred population presented an excess of homozygosity relative to controls. Next, we delineated the scale of inbreeding in AD cases. To prioritize regions with potential recessive loci, we constructed a homozygosity map of genomic regions overrepresented in inbred AD cases. Finally, we performed further exploration of several promising candidate ROHs using whole exome sequencing (WES) data.

## Subjects and Methods

The overview of the proposed strategy for ROH detection and subsequent prioritization is depicted in Figure 1.

**Figure 1.**
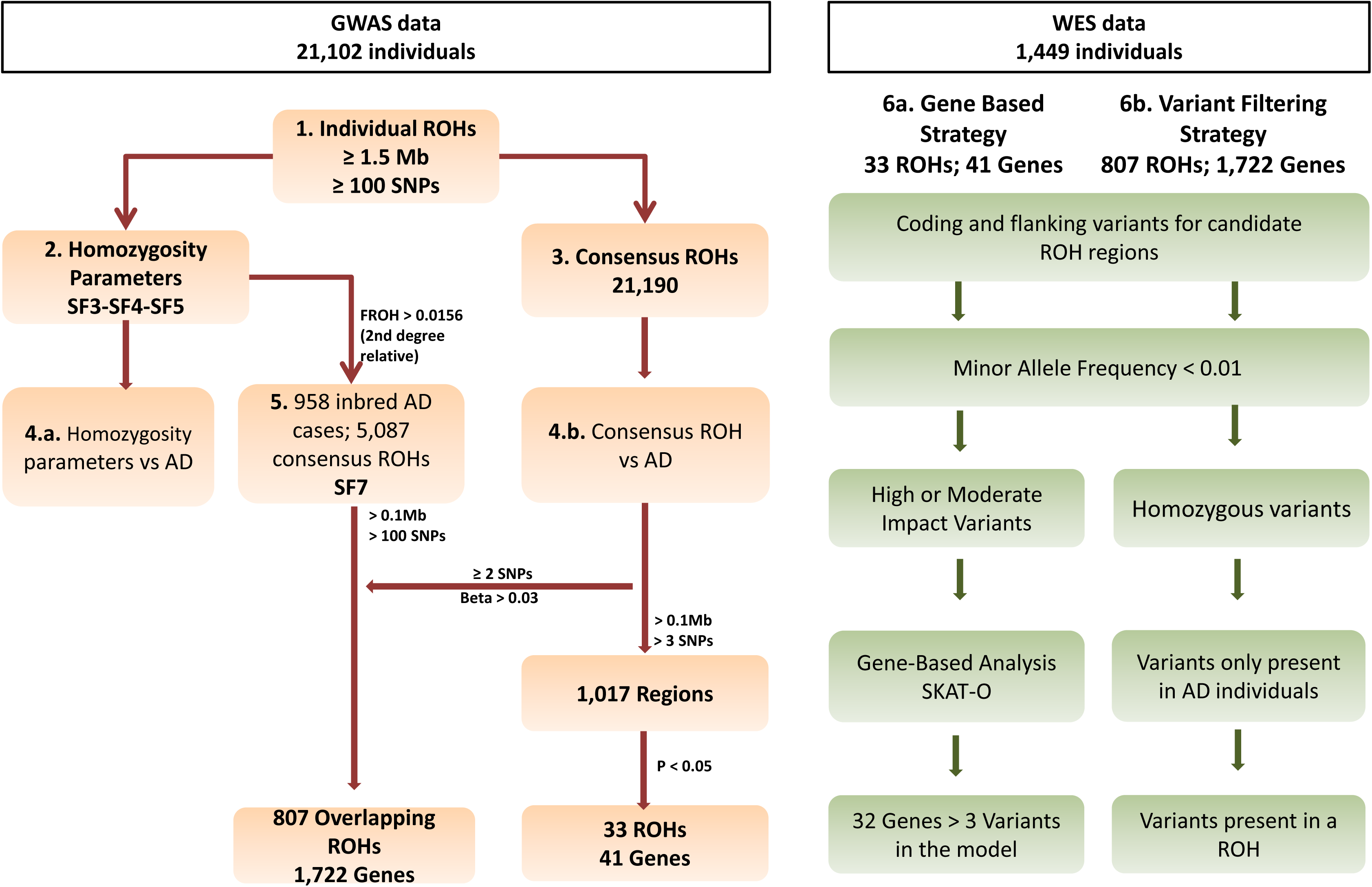
Schematic of the stepwise for ROH prioritization.

### Genotyping data

This study includes 10 independent genome-wide data sets comprising a total sample of 21,100 unrelated individuals (11,921 AD cases and 9,181 individual controls) of European ancestry (Supplementary Table 1). The recruitment, phenotyping, and quality control for genome-wide data has been described previously ^13^.

Briefly, genotype-level data for each cohort was processed by applying identical quality control and imputation procedures. Individuals were excluded for low-quality samples, (call rate <97%), excess heterozygosity, sample duplicates, or relation to another sample (PIHAT > 0.1875). Individuals were excluded if sex discrepancy was detected. Population outliers of European ancestry were also removed. Variants were excluded if they departed from the Hardy-Weinberg equilibrium (P-value ≤ 1 × 10^-6^), presented a different missing rate between cases and controls (P-value < 5 × 10^-4^ for the difference), or had a low frequency (MAF < 0.01) or low call rate < 95%. High-quality variants were imputed in Michigan Server using the Haplotype reference consortium (HRC) panel (https://imputationserver.sph.umich.edu). Only common markers (MAF > 0.05) with a high imputation quality (*R*^2^>0·90) were used for downstream analysis.

Next, we generated a merged data set combining imputed genotypes from available data sets. We calculated identity-by-descendent (IBD) with PLINK 1.9 to generate a cohort of unrelated individuals. All possible pairs had Pi-hat < 0.1875, a Z0 ≥ 0.75 and a Z1 ≤ 0.25. Imputed markers with call rates > 0.95 and MAF > 0.05 in the merged data set were selected for ROH calling (N _SNPs_ = 2,678,325).

### Whole Exome Sequence (WES) data

To meet the objective of exploring most promising ROH candidates in the sequencing data, we used the Knight-ADRC-NIA-LOAD (KANL) cohort ^33^. We excluded autosomal dominant familial cases and sporadic AD cases harboring well-known disease-causing mutations, as they could explain disease status. Thus, this study comprised 986 AD cases and 463 control individuals of European ancestry (See Supplementary Table 1 and Supplementary Figure 1). Of these, 488 subjects presented both GWAS and WES data available for this study. Detailed descriptions of cohort characteristics and quality control for WES data have been provided previously ^33^.

Briefly, exome libraries were prepared using Agilent’s SureSelect Human All Exon kits V3 and V5 or Roche VCRome. WES samples were sequenced on a HiSeq2000 with paired ends reads, with a mean depth of coverage of 50x to 150x for WES and 30x for WGS. Fastq sequences were aligned to the GRCh37.p13 genome reference. Variant calling was performed following GATKv.3.6 Best Practices (https://software.broadinstitute.org/gatk/) and restricted to a capture region with 100bp of padding. Variants and indels within 99.9% of the VQSR confidence interval were included in the analysis, along with variants with an allele balance between 0.30– 0.70, a quality depth ≥5 for indels and ≥ 2 for SNPs, and a call rate > 95% ^33^.

### 1- Identification of individual ROHs

Individual ROH calling was conducted using the observational genotype-counting approach implemented in PLINK (v1.09) (https://www.cog-genomics.org/plink/1.9/), as it outperforms additional methods in ROH detection ^34^. ROH detection was performed for each individual study and for the merged data set using imputed genotypes. Since included data sets were genotyped using different genotyping arrays, they shared a small fraction of directly genotyped markers. Given that it has been demonstrated that lower SNP density can impact the accuracy of ROH analysis ^35^, we decided to use high-quality imputed genotypes to increase SNP coverage. We used a sliding window of 50 SNPs of 5000 Kb in length to scan the genome. One heterozygote and five missing calls per window were tolerated in order to manage genomic regions with a small number of genotyping errors and discrete missingness. These parameters were similar to those described previously ^36^. The minimal number of SNPs in a ROH was set to 100 SNPs ^37, 38^. We empirically explored two minimal length cut-offs to consider a ROH, 1 Mb and 1.5 Mb. It has been suggested that ROHs > 1Mb prevent the detection of short homozygosity stretches, which, according to empirical studies ^39, 40, 41^, are generated by linkage disequilibrium forces in the human genome. However, the ability to detect autozygous regions with ROH length set to 1Mb could be compromised. Inbreeding estimations resulting from individual ROHs ≥ 1.5Mb have been most strongly correlated with inbreeding estimated from pedigree information ^23^, but this threshold has never been applied to AD studies. Autosomal SNPs were included in a ROH if >5% of the sliding window was homozygous. This means that at least 3 SNPs in 250 Kb from the sliding window were required to include a new marker. The maximum distance between two consecutive SNPs was set to 1000 Kb apart, and SNP density to at least 1 SNP in 50Kb.

### 2- Exploration of homozygosity parameters

To assess the data quality and genetic architecture of detected ROHs (> 1 Mb and > 1.5 Mb) in each individual study and in the whole dataset, we calculated: a) the mean of the total length of ROH or sum of ROH (SROH); b) the average ROH length (AVROH); c) the number of ROHs (NROH); and d) ROH-based estimates of the inbreeding coefficient, F, (FROH) per individual. AVROH is the SROH divided by NROH per subject. FROH represents the proportion of homozygous segments in the autosomal genome per individual (Equation 1). For individuals, this would be the SROH detected divided by a factor of 3,020,190 Kb, the total autosomal genome length according to the GRCh37.p13 assembly. We further explored whether the effect of homozygosity parameters was similar when: 1) ROH length was set to 1 Mb or 1.5 Mb; and 2) the analysis was performed per data set or in the final merged database (Supplementary Figure 2). Supplementary Table 2 and Supplementary Figure 3 demonstrate that FROH estimates derived from ROH calling at 1Mb exhibited a large degree of inflation, not allowing an accurate detection of inbreeding (Mean FROH 1Mb = 0.028; Mean FROH 1.5Mb = 0.011), which is in accordance with prior studies ^23^. After conducting an analysis of the 2,678,325 SNPs shared between available data sets, we found that the parameters of the individual data sets and the merged data set analyses were similar (Supplementary Figure 2 and Supplementary Table 3). After these exploratory analyses, we decided to conduct downstream analyses with ROH calling at 1.5 Mb in the merged data.

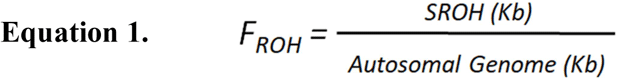

Copy number variants (CNV), particularly hemizygous deletions, are known to cause spurious ROHs. However, prior studies have demonstrated that the impact of performing ROH calling with or without CNVs is only 0.3% of the total ROH length ^23^, making it highly unlikely that deletions called as ROHs influence findings. To assess the impact of CNVs, specifically deletions, in our study, we also conducted ROH calling after removing common CNV deletions extracted from the Database of Genomic Variants (DGV) (http://dgv.tcag.ca/) ^42^.

### 3- Identification of consensus ROHs

Consensus ROHs were defined as overlapping segments between individual ROHs observed in different genomes. A consensus ROH needs a DNA segment match of at least 95% for non-missing SNP markers. Consensus ROH calling was performed using PLINK 1.9 in the merged data set. We then extracted those consensus ROHs with a DNA length over 100 Kb and more than 3 consecutive SNPs. These criteria were applied to prevent the detection of false positives.

### 4- Analyses

#### 4a-Association analysis between homozygosity parameters and AD risk

To assess the quality of the data in each individual study, we explored sample distribution for each of four homozygosity parameters: NROH, SROH, AVROH, and FROH. Exploratory analysis was depicted with violin plots, which combine a box plot with a kernel density plot, using the ggplot2 package from R (Supplementary Figure 4 and 5). Inverse rank normal transformation was performed to generalize homozygosity parameters using “rankNorm” option in the RNOmni package in R. Transformed distributions are shown in Supplementary Figure 6. To test the association of homozygosity parameters with AD status, we developed a generalized linear model for a binominal outcome, using R for individual-level data. To account for potential heterogeneity between individual studies, we adjusted the model per cohort and the first four principal components (PCs) resulting from ancestry analysis. See Equation 2. Sensitivity analysis was conducted to explore the impact of age on homozygosity parameters (Supplementary Table 6).

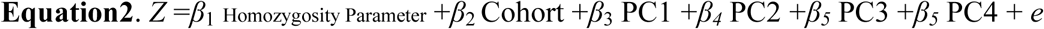

#### 4b-Association analysis between consensus ROHs and AD

The association between the phenotype and all consensus ROHs was explored using a logistic model. The model was adjusted per cohort, with PCs as covariates for downstream analysis. Nonetheless, covariate models adjusted for age and gender, in addition to cohort and PCs, were also calculated. Regression-based results were corrected for multiple testing using a Bonferroni correction.

Next, we sought to estimate whether there was an overrepresentation of risk (β > 0) or protective (β < 0) consensus ROHs in our association results at different levels of length and SNP number per consensus ROH. We applied a binominal test using R. We considered that under the null hypothesis of no association, similar distributions would be expected for both (50/50).

### 5- The homozygosity map of inbred AD individuals

#### 5a-Identificationn of inbred individuals

We used FROH estimates to detect the subset of inbred individuals within the whole data set. This parameter has been previously shown to better correlate with the unobserved pedigree inbreeding ^32, 43^. The cut-off between inbred and non-inbred individuals was set to F_ROH_ > 0.0156 ^35^. This cut-off corresponds to a second-degree relation, i.e. the mean inbreeding coefficient for kinship in a second-cousin marriage or closer. It was assumed that there are no different biological effects below 0.0156 than in the general population ^44^. We demonstrated the efficient capture of inbred individuals as indicated in Supplementary Figure 7, which shows the inverse relationship between ROH length and ROH age. Thus, short ROHs evidence ancient origin, and long ROHs more recent origin, which might indicate ROHs emerging from consanguineous mating. Next, to explore whether the frequency of consanguinity was higher in AD cases than in controls, we calculated the odds ratio and chi square p values using the epitools package in R.

#### 5b-ROHs prioritization based on inbred AD cases

ROH detection was conducted in the subset of inbred AD cases, applying similar criteria to those previously described for the outbred population. Briefly, considering the long size of homozygous tracts for inbred individuals, there is higher probability of finding a consensus ROH by chance within consanguineous AD cases than in the general population. Hence, we applied stringent criteria to define consensus ROHs. Consensus ROHs from inbred AD cases with ROH lengths > 100 Kb and ROH > 100 SNPs were given priority for further analysis. Shared overlapping regions between inbred AD cases and the whole data set were also identified (See bash code in Supplementary Material), and selected based on their overrepresentation in AD cases relative to controls (β > 0.03). Prioritized regions were then explored in sequencing data.

### 6- Candidate gene prioritization strategies using WES

#### 6a-Gene based analysis

To prioritize genes in consensus ROH regions, we performed a gene-based analysis (986 cases vs 463 controls) (Figure 1). To generate SNP sets, variants were filtered out according to minor allele frequency (MAF<0.01) and functional impact. The allele frequency cut-off was established according the Exome Aggregation Consortium (ExAC), non-Finnish European Exome Sequencing project (ESP), and 1000G. Only those variants predicted to have a high or moderate effect according to SnpEff were included ^45^. To compute p-values per gene set, SKAT-O model were applied using R. The models were adjusted to consider the impact of the first two PCs and sex. Genes were filtered out from results if the number of SNPs included in the model was less than or equal to 3.

#### 6b-Variant filtering strategy for inbred AD cases

ROH segments emerging from inbred AD cases are the most promising candidates to harbor autosomal recessive variants. Therefore, we deeply explored ROHs by applying an alternative strategy based on variant filtering. In the present study, we explored 488 AD cases with complementary GWAS and WES data to identify candidate genes and/or mutations associated with AD. Because there is a low likelihood to identify any novel or causative mutation in available databases, variants with MAF > 0.01 in the Exome Aggregation Consortium (ExAC), non-Finnish European Exome Sequencing project (ESP), and 1000G were excluded. All heterozygous variants were removed. Finally, only the variants mapped in individual ROHs were selected.

#### Biological significance of ROH findings

To map genes within ROHs, we first extracted all the SNPs located in ROH regions. Next, we individually annotated each one.

## Results

### ROH parameters are associated with Alzheimer’s disease risk

We examined the general characteristics of the four ROH parameters (SROH, NROH, AVROH, FROH) in 21,100 unrelated European individuals from 10 independent cohorts (Supplementary Table 1). Data distributions in each individual data set and in the joint analysis are shown in Supplementary Table 2 and Supplementary Figure 4. Relationships between the mean NROH and SROH are shown in Figure 2. Within the merged data set the mean NROH was 14.6 ± 4.6, the AVROH was 2.11 ± 0.61Mb, and the SROH was 31.9 ± 22.2Mb. These estimations are in accordance with those observed in European individuals ^35^, except for the NROH parameter, which was higher than in previous studies ^35^.

**Figure 2.**
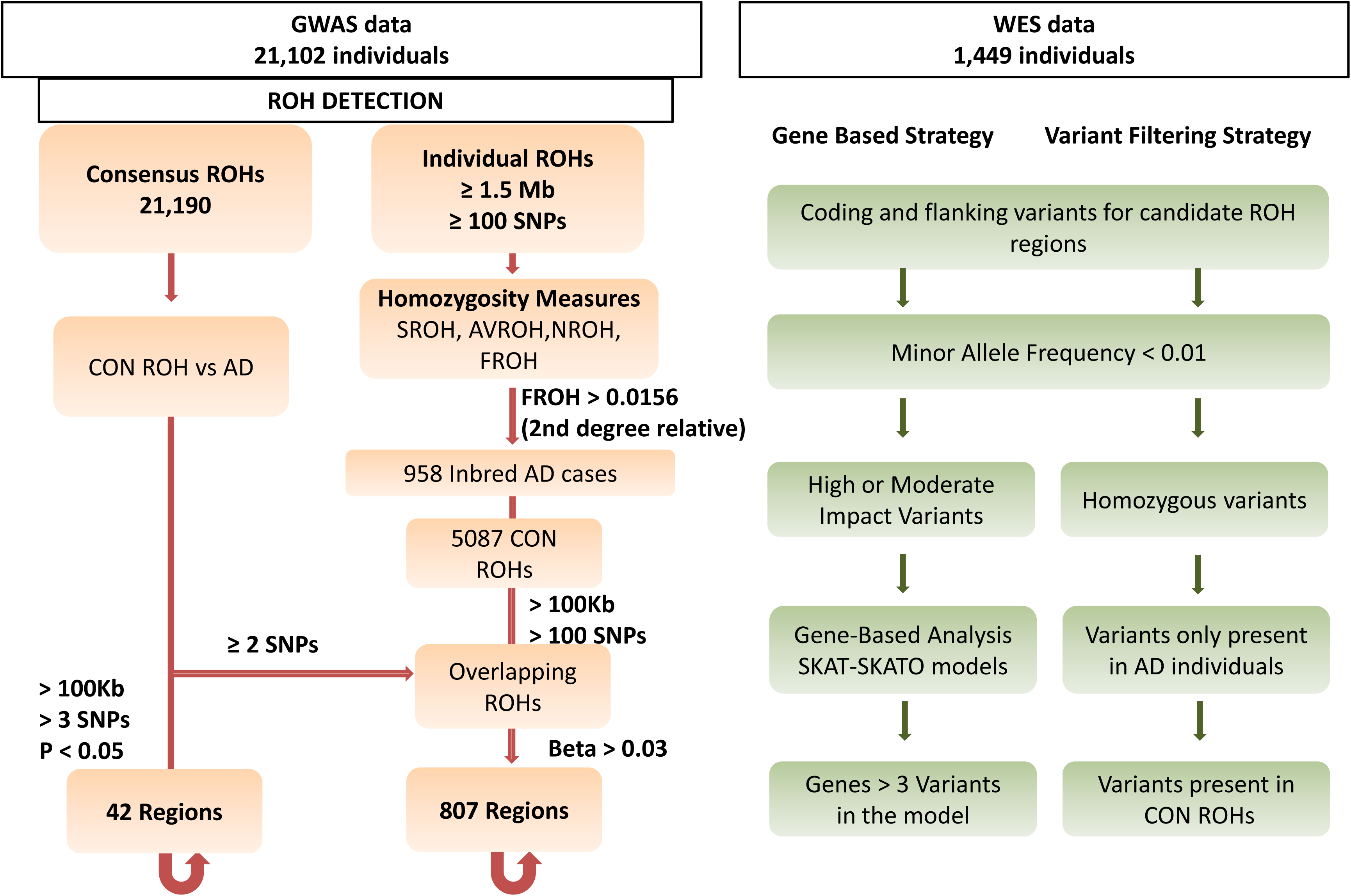
Runs of homozygosity per cohort and per individual. A) Mean number of ROHs versus mean total sum of ROHs in Mb for the 10 cohorts explored. B) Mean number of ROHs versus mean total sum of ROHs in Mb per individual explored. Red dashed lines represent the threshold for the inbreeding coefficient of 0.0156 (second cousins’ offspring) and 0.0625 (first cousins’ offspring).

Next, we tested the association of the four parameters between AD cases and control subjects. We found that i) a larger homozygosity fraction of the genome (F_ROH_) increased the risk of suffering AD [β _FROH_ (CI95%) = 0.051 (0.023 – 0.078); p value = 3.25 x 10^-4^] (Table 1); ii) AD individuals presented more ROH segments compared to controls [β _FROH_ (CI95%) = 0.043 (0.015 – 0.071); p value = 2.48 x 10^-3^]; iii) and average lengths of ROHs were increased in AD cases compared with controls [β _FROH_ (CI95%) = 0.027 (0.000 – 0.055); p value = 0.051] (Table 1). Results for each cohort are shown in Supplementary Table 4. Notably, a sensitivity analysis conducted excluding known deletions, i.e. hemizygous segments ^42^, provided similar results (Supplementary Table 5).

**Table 1.** Effect of genome-wide homozygosity measures in Alzheimer’s disease for the joint analysis.

| Dataset | Unadjusted |  | Adjusted Cohort |  | Adjusted Cohort, PCs |  |
| --- | --- | --- | --- | --- | --- | --- |
|  | Beta (CI95%) | P value | Beta (CI95%) | P value | Beta (CI95%) | P value |
| **FROH** | 0.059<br>(0.032 - 0.087) | 2.02 x<br>10<sup>-5</sup> | 0.064<br>(0.036 - 0.091) | 4.96 x<br>10<sup>-6</sup> | 0.051<br>(0.023 - 0.078) | 3.25 x<br>10<sup>-4</sup> |
| **AVROH** | 0.014<br>(-0.001 - 0.053) | 0.060 | 0.030<br>(0.003 - 0.057) | 0.032 | 0.027<br>(0.000 - 0.055) | 0.051 |
| **NROH** | 0.053<br>(0.027 - 0.081) | 1.11 x<br>10<sup>-4</sup> | 0.058<br>(0.030 - 0.086) | 3.62 x<br>10<sup>-5</sup> | 0.043<br>(0.015 - 0.071) | 2.48 x<br>10<sup>-3</sup> |

*Results for association of excess of homozygosity ( $F_{ROH}$ ), average ROH length (AVROH), and number of ROH (NROH) with Alzheimer disease status.*
*OR, Odds Ratio; with 95% confidence interval (CI95%) and level of statistical significance (P value)*

We also detected a correlation between age and homozygosity measures in the control group populations ranging from 50 to 80 years old. Specifically, FROH and NROH exhibited a significant positive correlation. Conversely, AVROH showed a significant inverse correlation with age (Supplementary Table 6). Given these findings, we decided to test the impact of acquired clonal mosaicism introduced by aging on homozygosity estimations. First, we conducted a sensitivity analysis, controlling for cohort, PCs, and age. The effect of FROH remained significant and stable after adjustments. The average length of ROH also remained significantly different between cases and controls [β _AVROH_ (CI95%) = 0.074 (0.040 - 0.106); p value = 2.16 x 10^-5^)]. Interestingly, the number of ROHs was largely age-dependent (p value = 8.93 x 10^-9^), and it was not significantly associated with AD after age adjustment [(β _NROH_ (CI95%) = 0.010 (−0.024 - 0.044); p value = 0.559]. These findings support the notion that genomic somatic instability increases with age and can pervasively distort the gene-dosage of multiple loci (Supplementary Table 6).

### ROH analysis of AD risk using the whole data set

We identified 21,190 consensus ROHs in the merged data set (*N* = 21,100). We then tested the association of each consensus ROH with AD status. Of these, 11,974 were found to be enriched in AD cases, and 9,216 were enriched in controls. Overall, we observed a highly significant over-representation of ROHs that increased the risk of suffering AD (p value < 2.20 x 10^-16^) (Table 2). The same over-representation of risk associations was detected after filtering at several levels based on the length and number of SNPs per consensus ROH (Table 2). When the test was conducted with results adjusted for cohort, PCs, age, and gender, the over-representation of risk associations remained significant (p value < 2.20 x 10^-16^).

**Table 2.** Frequency of consensus ROHs with a potential risk or protective effect in Alzheimer’s disease.

|  | **N**<br>**ROH** | **Risk**<br>**associations** | **Protective**<br>**associations** | **P value** | **Probability of**<br>**Success** |
| --- | --- | --- | --- | --- | --- |
| **Whole dataset** | 21190 | 11974 | 9216 | $< 2.2 \times 10^{-16}$ | 0.56 |
| **Category A** | 1017 | 593 | 424 | $< 2.2 \times 10^{-16}$ | 0.58 |
| **Category B** | 926 | 537 | 389 | $1.30 \times 10^{-6}$ | 0.57 |
| **Category C** | 858 | 499 | 359 | $1.98 \times 10^{-6}$ | 0.58 |
| **Category D** | 42 | 33 | 9 | $2.7 \times 10^{-4}$ | 0.79 |
| **Whole dataset /**<br>**Map of Inbreed AD**<br>**Cases** | 6636 | 3969 | 2667 | $< 2.2 \times 10^{-16}$ | 0.60 |

*Strategy A, ROHs > 100 kb; > 3 SNPs*
*Strategy B, ROHs > 100 kb; > 25 SNPs*
*Strategy C, ROHs > 100 kb; > 50 SNPs*
*Strategy D, ROHs > 100 kb; > 3 SNPs, $P < 0.05$*

To prevent the detection of false positive associations, we selected consensus ROHs with ≥100 Kb and ≥3 SNPs, which provided a subset of 1,017 consensus ROHs (Figure 1 and Supplementary Table 7). After correction of multiple tests (Bonferroni correction of p = 4.92 x 10^-5^), the most significantly associated ROH was detected in 57 individuals (45 AD cases vs 12 controls, β (CI95%) = 1.09 (0.48 ‒ 1.48), p value = 9.03 x 10^-4^). It expanded 115.9Kb into an intergenic region (chr4:11,189,482‒11,305,456) near the *HS3ST1* locus. This region survived age and gender adjustments (Supplementary Table 7). Importantly, this region has been previously associated with AD ^46^, but the recessive model has never been tested.

Using associated ROH as a reference, we explored the genes located in significant risk consensus ROHs (p value < 0.05) in WES data as well (Figure 1). A total of 33 ROHs comprising 41 genes were analyzed. Of those genes, 32 included >3 SNPs in the model (32 genes; Bonferroni correction p value = 0.0015). The *NECAB1* locus (chr8:91,803,921-91,971,630) presented the most significant signal (p = 0.01) (Supplementary Table 8), but none loci reached the multiple test correction threshold.

### Homozygosity mapping of AD using DNA segments identified in inbred cases

We detected 1,621 individuals presenting a F_ROH_ ≥ 0.0156 among the total sample (N = 21,100) (Figure 2) (Supplementary Table 9). Interestingly, inbreeding over the second degree of consanguinity was associated with higher risk of suffering AD [OR (95%, CI) = 1.12 (1.01 – 1.25); p value = 0.027), which is in line with our previous results. This supports the idea that consanguineous AD cases are overrepresented in the general AD population. Accordingly, the search for recessive loci that play a role in AD can first be assessed in consanguineous cases.

After ROH calling in inbred AD cases, we detected 5,087 pools of overlapping ROHs. From these, we extracted consensus ROHs with ≥100 Kb and ≥100 SNPs. We then selected those ROHs that overlapped with the whole sample and that were over-represented in AD cases relative to controls (Figure 1). We prioritized 807 consensus homozygous segments from inbred cases (Figure 3 and Supplementary Table 10). Together, these represented 8.6% of the total autosomal genome and comprised 1,722 genes (Supplementary Table 11). Of these, 1,136 genes, including >3 SNPs in the model, were further explored in WES data using a gene-based approach. None of them remained associated after multiple corrections (N _genes tested_ = 1,136; p value = 3.47 x 10^-5^). Our top signal was detected in the *FRY* locus (p value = 0.001) (Supplementary Table 11).

**Figure 3.**
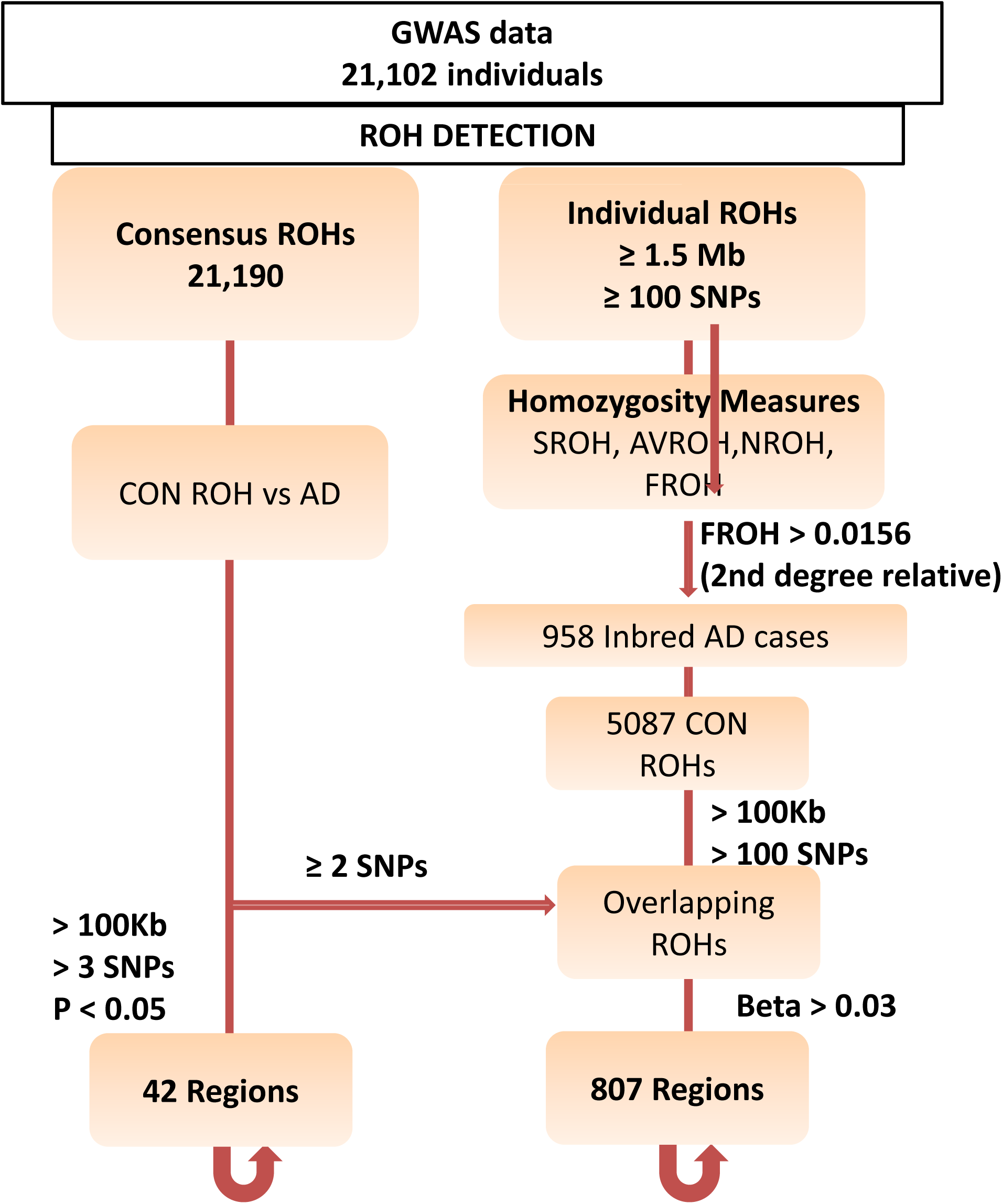
Circos plot for the prioritized regions. Histogram for the effect of the 21,190 consensus ROHs identified in the whole sample is shown. Risk ROH associations are shown in red; protective ROH associations are shown in green. Blue regions represent prioritized ROHs from consanguineous AD cases. Orange segments represent prioritized regions harboring potential recessive variants.

Considering that recessive variants are expected at low frequencies, even gene-based analysis would be underpowered to detect significant associations. Therefore, we decided to further prioritize loci by searching homozygous mutations within selected consensus ROHs from inbred AD subjects (Figure 1). We identified seven AD cases that had eight new (or extremely rare) homozygous variants in long ROH segments (Table 3). Two of these individuals were consanguineous (F _ROH_ > 0.156). One had a missense variant (rs140790046, c.926A>G) that encodes p.Asn309Ser change within the *MKX* locus. Another carried a novel variant (rs116644203) in the *ZNF282* locus, which was located in an extremely large region of homozygosity (14.9 Mb) (Table 3). Furthermore, three additional homozygous variants were detected: i) a variant (rs117458494) in the *SPON1* locus, previously related with amyloid metabolism ^47^, and ii) two potential causative variants, carried only by this individual, within a previously identified AD region (*TP53INP/NDUFAF6)* ^13^. One (rs73263258-*ESRP1*; in *TP53INP/NDUFAF6* region) is a missense variant (c.475G>A) that encodes p.Ala159Thr change (Table 3). Further notes and functional effect predictions for these variants are provided in Supplementary Table 12.

**Table 3.** Candidate recessive variants after ROH prioritization focused on inbred AD cases.

| Individual | Froh<br>ROH>1500kb | SROH<br>(Kb) | CH<br>R | ROH start | ROH end | ROH<br>length | ROH<br>SNPs | Rs | Variant | Near<br>Locus | Ref<br>Allele | Alt<br>Allele | MAF | Impact |
| --- | --- | --- | --- | --- | --- | --- | --- | --- | --- | --- | --- | --- | --- | --- |
| Individual<br>1 | 0.0058 | 17421.6 | 7 | 53232690 | 55262627 | 2029.938 | 4213 | NA | 7:53930748 | RP11-678B3.2 | C | A | Novel | MODIFIER |
| Individual<br>2 | 0.0185 | 55861.4 | 7 | 13561771<br>5 | 15051839<br>3 | 14900.679 | 12426 | rs116644203 | 7:148903805 | ZNF282 | C | T | Novel | LOW<br>MODIFIER |
| Individual<br>3 | 0.0099 | 29868.8 | 8 | 47018484 | 49544784 | 2526.301 | 1265 | NA | 8:48352954 | SPIDR | T | A | 0.004 |  |
| Individual<br>4 | 0.0153 | 46194.5 | 8 | 91282507 | 96735572 | 5453.066 | 4734 | rs73263258 | 8:95658495 | ESPR1* | G | A | 0.002 | MODERAT<br>E<br>MODIFIER |
|  |  |  |  |  |  |  |  | rs138998679 | 8:95780656 | DPY19L4* | A | G | 0.003 | LOW |
| Individual<br>5 | 0.0179 | 54137.7 | 10 | 24516308 | 28414933 | 3898.626 | 6248 | rs140790046 | 10:27964291 | MKX | T | C | 0.0005 | MODERAT<br>E |
| Individual<br>6 | 0.0137 | 41392.6 | 11 | 12032272 | 16358722 | 4326.451 | 4473 | rs117458494 | 11:14282085 | SPON1<br>RP11-21L19.1 | G | A | Novel | MODIFIER |
| Individual<br>7 | 0.0115 | 34603.4 | 14 | 63906601 | 66333670 | 2427.07 | 2633 | rs140897155 | 14:64688393 | SYNE2<br>ESR2 | G | A | 0.001 | MODERAT<br>E<br>LOW<br>MODIFIER |

## Discussion

This study represents the largest analysis of homozygosity (ROHs) conducted in the context of Alzheimer’s disease (AD). Our estimate of excessive homozygosity in individuals with AD from European populations (N = 21,100) provides firm evidence for the role of consanguinity in AD. This finding suggests that there might be numerous recessive AD loci. This statement has several implications for the design of AD genetic studies, for the better understanding of the causes of phenotypic variation in AD and, finally, for the search for an efficient therapeutic target.

In this study, we efficiently identified numerous potentially inbred AD cases nested in an outbred population, offering a new framework for the analysis of the inbreeding component in AD. Furthermore, we demonstrated that detected consensus ROHs are enriched in risk associations. Considering our findings, we believe that recessive allelic architecture defines a portion of AD heritability. Even the accumulation of multiple low-penetrance or pure recessive variants is likely to play a role.

The genetic basis of human late-onset diseases has been mainly explained by selective neutrality ^48^ under the common disease-common variant (CD-CV) hypothesis ^49^. Considering that, several evolutionary theories of aging have been proposed and demonstrated to some extent, e.g. mutation accumulation (MA) and antagonistic pleiotropy (AP) ^50, 51^. Theoretical and experimental models for MA further support inbreeding effects for late-onset diseases ^52, 53^. Given that the human population is evolutionarily young, a large degree of human variation is necessarily rare ^54^. Thereby, late-acting alleles will also be found at low frequencies. In that scenario, contemporary genetic models are in agreement with present results, where rare and recessive-acting variants could explain a part of the genetic basis of AD.

Previous studies in populations from European and non-European ancestries have shown inconsistent results ^26, 27, 28, 29^ in searches for homozygosity patterns in AD. We believe that several technical considerations must be taken into account in the analysis of ROH. First, it is suggested that the estimation of an excess of homozygosity in an outbred population requires a large sample size ^32^, but prior studies had very modest sample sizes (N < 6,000) ^26, 27, 28, 29^. Thus, they likely were underpowered to detect inbreeding effects from unrelated individuals.

Second, different scenarios should be considered for selecting shorter or longer ROHs than 1.5 Mb for the measurements and statistics, because these indicate different aspects of demographic history. Evidence suggests that individual ROHs < 1.5 Mb might reflect LD patterns of ancient origin rather than the consanguineous cultural practices and genetic isolation captured with ROHs > 1.5 Mb ^23^. Here, we detected substantial inflation in FROH estimations when individual ROH length was set to 1 Mb. This makes the detection of inbred individuals from an outbred population complicated, and strongly confounds the interpretation of homozygosity estimations. Despite that, prior AD genetic studies assessing the role of homozygosity have not tested the potential effect of performing ROH calling for segments longer than 1 Mb ^26, 27, 28, 29^, which might partly explain initial failures.

Overall, several technical handicaps have made difficult ROH studies in AD. It might have caused researchers to overlook the potential inbreeding effect for this disease. Hence, we encourage other groups to conduct ROH analysis in unrelated populations, but with large enough sample sizes and while redefining the ROH lengths at least to 1.5Mb, to better capture the recessive component of AD.

In our ROH analysis using the whole sample (N = 21,100), the most promising consensus ROH was located in proximity to the *HS3ST1* gene (⁓200 Kb), and showed a strong genetic effect. Genetic markers in the vicinity of the homozygous block (⁓300kb) have been previously associated with AD ^46, 55^. Additionally, the *HS3ST1* locus was differentially expressed in the brain in AD cases versus controls ^55^. Despite these findings, the causative genetic mechanism involving this region with AD remains elusive. We therefore believe that high-resolution mapping across the 115 Kb of the reported consensus ROH could help to positional cloning of the causative mutation.

Our study of ROHs > 1.5Mb sheds light on the homozygosity component influencing AD, as it reflects recent consanguinity and/or population isolation. Inbred individuals tend to have lower survival, fertility, and growth rates ^56, 57, 58^, as well as post-reproductive health ^59^. Considering that, we believe that enriching our subset of inbred cases can provide a redefined framework for investigating inbreeding effects and looking for recessive acting variants. This idea has driven the design of the present study. With the aim to increase the probability to detect regions harboring recessive-acting loci, we prioritized consensus ROHs according to the homozygosity map of inbred AD individuals that we obtained. Candidate regions were then explored in sequencing data. Among them, variants of the *MKX* and *ZNF282* genes were detected in two independent inbred AD cases. Both *ZNF282* and *MKX* loci are encoding transcription factors ^60, 61, 62^. This is worth noting, given that the largest WES study analyzing rare variations in AD recently highlighted the potential role of transcriptional regulation for this disease ^63^. As well, the *ZNF282* gene is mapped roughly to 800Kb from the *CNTNAP2* gene, which has been previously associated with AD ^11^. Autosomal recessive mutations in *CNTNAP2* loci have been also linked with epilepsy and intellectual disability (<u>OMIM 604569</u>).

In this study, we also found a potential recessive variant in the *SPON1* locus. SPON1 has been related to the mechanism of AD, where APP metabolism has a central role. APP cleavage through β-secretases produces amyloid-beta (Aβ), which later accumulates in AD brains ^7^. SPON1 has been found to bind to APP, inhibiting its α/β cleavage ^47^. Other studies have also reported SPON1 binding to the APOE family of receptors ^64^. Markers in this gene have been related with dementia severity ^65^ and with the rate of cognitive decline ^66^. Considering prior findings and the present result, it would be biologically plausible that the presence of recessive acting-variants in APP or in its biological partners directly influences the amyloid cascade. Thus, we believe that *SPON1* could be considered an interesting candidate, which deserves future resequencing efforts.

Among other candidates, we identified a missense variant (rs73263258 in *ESPR1* gene) within a long ROH in an AD patient. This gene was mapped in the close vicinity of the *TP53INP1/NDUFAF6* genomic region. This region has been previously associated with AD using a gene-based strategy ^67^. Recently, our group also identified genome-wide significant markers in this region ^13^. It is not unexpected that genes containing common variants with small genetic effect might also be enriched in rarer variants with higher penetrance. The existence of several genetic mechanisms acting in this region should be considered when deep sequencing will be conducted, to help pinpoint the causative variant.

Our observations are subject to limitations that need to be considered. Since the data sets used in the present study were genotyped using different platforms, they shared a small proportion of directly genotyped markers. Given that lower SNP density could impact the accuracy of the study ^35^, we decided to perform the present analysis using imputed genotypes of high quality (imputation quality, *r*2> 0.90). To make the data optimally comparable, we generated a merged data set including the same variants with MAF > 0.05. We also showed that ROH calling is insensitive to performances of the analysis in each individual dataset or in the merged data for a set of individuals from the same ancestral group, when we determined the SNP set to use.

The potential impact of CNVs on ROH analysis must be taken in consideration as a potential limitation. However, when we assessed CNV impact on our analyses, no differences were found in homozygosity parameters before and after CNV exclusion. These results are in agreement with those of previous studies, suggesting that the effect of deletions on homozygosity parameters, when it exists, is minimal ^36^.

Clonal mosaicism can also generate spurious ROHs. A direct correlation between clonal mosaicism events in peripheral blood and age >50 years was demonstrated ^68^. We believe that these events might be introducing ROHs of short lengths. Consequently, an age-dependent increase in the number of segments was detected. In fact, it could explain why we identified a higher-than-expected mean ROH number for this data set of a European population than was found in prior studies ^35^. From our point of view, controlling the role of genome instability for late-onset neurodegenerative diseases represents a challenge, due to the impossibility of detecting true converters to AD before disease onset, and the difficulties of collecting biological information from the target tissue. Despite the fact a signature of genome instability in ROH studies might exist, our adjusted results (those considering age effects) still support the idea that inbred individuals are overrepresented in AD population respect to controls.

In summary, we demonstrated the existence of an inbreeding effect in AD and efficiently captured a fraction of consanguineous individuals from outbred populations. The proposed method can be considered a refined strategy to investigate the role of recessive variants in AD. Considering that there are significant barriers to collecting complete information from consanguineous AD families, the identification of highly probable consanguineous AD cases in outbred populations could be important for future large-scale homozygosity mapping. Furthermore, the opportunity to explore complementary sequencing data gave an added value to this research, providing a subset of potential candidates harboring recessive variants. In any case, the proposed candidates, acting under a recessive inheritance model, will only be confirmed when at least an additional individual harboring the same recessive mutation, or a compound heterozygote is detected. We recognize our current lack of power to fully verify arAD loci. That is why greater efforts and larger collections of individuals with GWAS and sequencing data are needed to confirm our findings.

Our understanding of the dynamics of population genomics in complex diseases like AD is far from complete, but ROH analyses provide us a means to go further and might be an alternative strategy to uncover the genetic loci underlying Alzheimer’s disease.

## Acknowledgements

We would like to thank patients and controls who participated in this project. The Genome Research @ Fundació ACE project (GR@ACE) is supported by Fundación bancaria “La Caixa,” Grifols SA, Fundació ACE, and ISCIII (Ministry of Health, Spain). We also want to thank the private sponsors who support the basic and clinical projects of our institution (Piramal AG, Laboratorios Echevarne, Araclon Biotech S.A., and Fundació ACE). We are indebted to the Trinitat Port-Carbó legacy and her family for their support of Fundació ACE research programs. Fundació ACE is a participating center in the Dementia Genetics Spanish Consortium (DEGESCO). A.R. and M.B. receive support from the European Union/EFPIA Innovative Medicines Initiative Joint Undertaking ADAPTED and MOPEAD projects (Grants No. 115975 and 115985, respectively). M.B. and A.R. are also supported by national grants PI13/02434, PI16/01861, PI17/01474 and PI19/01301. Acción Estratégica en Salud is integrated into the Spanish National R + D + I Plan and funded by ISCIII (Instituto de Salud Carlos III)-Subdirección General de Evaluación and the Fondo Europeo de Desarrollo Regional (FEDER-“Una manera de Hacer Europa”). LMR is supported by Consejería de Salud de la Junta de Andalucía (Grant PI-0001/2017). Control samples and data from patients included in this study were provided in part by the National DNA Bank Carlos III (www.bancoadn.org, University of Salamanca, Spain) and Hospital Universitario Virgen de Valme (Sevilla, Spain); they were processed following standard operating procedures with the appropriate approval of the ethical and scientific committees of these institutions. The present work was performed as part of the Biochemistry, Molecular Biology and Biomedicine doctoral program of S. Moreno-Grau at Universitat Autònoma de Barcelona (Barcelona, Spain).

This work was supported by grants from the National Institutes of Health (R01AG044546, P01AG003991, RF1AG053303, R01AG058501, U01AG058922, RF1AG058501 and R01AG057777) and the Alzheimer’s Association (NIRG-11-200110, BAND-14-338165, AARG-16-441560 and BFG-15-362540). The recruitment and clinical characterization of research participants at Washington University were supported by NIH P50 AG05681, P01 AG03991, and P01 AG026276. This work was supported by access to equipment made possible by the Hope Center for Neurological Disorders, and the Departments of Neurology and Psychiatry at Washington University School of Medicine.

Data collection and sharing for this project was partially funded by the **Alzheimer’s Disease Neuroimaging Initiative (ADNI)** (National Institutes of Health Grant U01 AG024904) and DOD ADNI (Department of Defense award number W81XWH-12-2-0012). ADNI is funded by the National Institute on Aging and the National Institute of Biomedical Imaging and Bioengineering, as well as through generous contributions from the following: AbbVie; the Alzheimer’s Association; the Alzheimer’s Drug Discovery Foundation; Araclon Biotech; BioClinica, Inc.; Biogen; Bristol-Myers Squibb Company; CereSpir, Inc.; Cogstate; Eisai Inc.; Elan Pharmaceuticals, Inc.; Eli Lilly and Company; EuroImmun; F. Hoffmann-La Roche Ltd and its affiliated company Genentech, Inc.; Fujirebio; GE Healthcare; IXICO Ltd.; Janssen Alzheimer Immunotherapy Research & Development, LLC.; Johnson & Johnson Pharmaceutical Research & Development LLC.; Lumosity; Lundbeck; Merck & Co., Inc.; Meso Scale Diagnostics, LLC.; NeuroRx Research; Neurotrack Technologies; Novartis Pharmaceuticals Corporation; Pfizer Inc.; Piramal Imaging; Servier; Takeda Pharmaceutical Company; and Transition Therapeutics. The Canadian Institute of Health Research provides funds to support ADNI clinical sites in Canada. Private-sector contributions are facilitated by the Foundation for the National Institutes of Health (www.fnih.org). The grantee organization is the Northern California Institute for Research and Education, and the study was coordinated by the Alzheimer’s Therapeutic Research Institute at the University of Southern California. ADNI data are disseminated by the Laboratory for Neuro Imaging at the University of Southern California.

The **AddNeuroMed** data are from a public-private partnership supported by EFPIA companies and SMEs as part of InnoMed (Innovative Medicines in Europe), an integrated project funded by the European Union under the Sixth Framework program priority FP6-2004-LIFESCIHEALTH-5. Clinical leads responsible for data collection are Iwona Kłoszewska (Lodz), Simon Lovestone (London), Patrizia Mecocci (Perugia), Hilkka Soininen (Kuopio), Magda Tsolaki (Thessaloniki) and Bruno Vellas (Toulouse). Imaging leads are Andy Simmons (London), Lars-Olad Wahlund (Stockholm) and Christian Spenger (Zurich). Bioinformatics leads are Richard Dobson (London) and Stephen Newhouse (London).

Funding support for the **Alzheimer’s Disease Genetics Consortium (ADGC)** was provided through the NIA Division of Neuroscience (U01-AG032984).

The genotypic and associated phenotypic data used in the study “Multi-Site Collaborative Study for Genotype-Phenotype Associations in Alzheimer’s Disease (**GenADA**)” were provided by GlaxoSmithKline, R&D Limited. The data sets used for the analyses described in this manuscript were obtained from dbGaP at http://www.ncbi.nlm.nih.gov/gap through dbGaP accession number phs000219.

The **Mayo Clinic Alzheimer’s Disease Genetic Studies**, led by Dr. Nilüfer Ertekin-Taner and Dr. Steven G. Younkin at the Mayo Clinic in Jacksonville, FL, used samples from the Mayo Clinic Study of Aging, the Mayo Clinic Alzheimer’s Disease Research Center, and the Mayo Clinic Brain Bank. Data collection was supported through funding from NIA grants P50 AG016574, R01 AG032990, U01 AG046139, R01 AG018023, U01 AG006576, U01 AG006786, R01 AG025711, R01 AG017216 and R01 AG003949, NINDS grant R01 NS080820, the CurePSP Foundation, and the Mayo Foundation.

The **Neocodex-Murcia** study was funded by the Fundación Alzheimur (Murcia), the Ministerio de Educación y Ciencia (Gobierno de España), Corporación Tecnológica de Andalucía, Agencia IDEA (Consejería de Innovación, Junta de Andalucía), the Diabetes Research Laboratory, and the Biomedical Research Foundation. University Hospital Clínico San Carlos has been supported by CIBER de *Diabetes y Enfermedades Metabólicas Asociadas* (CIBERDEM); CIBERDEM is an ISCIII Project.

The **ROS/MAP** study data were provided by the Rush Alzheimer’s Disease Center, Rush University Medical Center, Chicago. Data collection was supported through funding from NIA grants P30AG10161, R01AG15819, R01AG17917, R01AG30146, R01AG36836, U01AG32984 and U01AG46152, the Illinois Department of Public Health, and the Translational Genomics Research Institute.

The **TGEN** study was supported by Kronos Life Sciences Laboratories, the National Institute on Aging (Arizona Alzheimer’s Disease Center grants P30 AG19610 and RO1 AG023193, the Mayo Clinic Alzheimer’s Disease Center grant P50 AG16574, and the Intramural Research Program), the National Alzheimer’s Coordinating Center (U01 AG016976) and the state of Arizona.

***The results published here are in part based on data obtained from the AMP-AD Knowledge Portal accessed at*** http://dx.doi.org/doi:10.7303/syn2580853

We thank the International Genomics of Alzheimer’s Project (IGAP) for providing summary results data for these analyses. The investigators within IGAP contributed to the design and implementation of IGAP and/or provided data but did not participate in analysis or writing of this report. IGAP was made possible by the generous participation of the control subjects, the patients, and their families. The i–Select chip was funded by the French National Foundation on Alzheimer’s Disease and Related Disorders. EADI was supported by the LABEX (laboratory of excellence program investment for the future) DISTALZ grant, Inserm, Institut Pasteur de Lille, Université de Lille 2, and the Lille University Hospital. GERAD was supported by the Medical Research Council (Grant n° 503480), Alzheimer’s Research UK (Grant n° 503176), the Wellcome Trust (Grant n° 082604/2/07/Z), and German Federal Ministry of Education and Research (BMBF): Competence Network Dementia (CND) grants 01GI0102, 01GI0711, and 01GI0420. CHARGE was partly supported by the NIH/NIA grant R01 AG033193 and the NIA AG081220 and AGES contract N01–AG–12100, the NHLBI grant R01 HL105756, the Icelandic Heart Association, and the Erasmus Medical Center and Erasmus University. ADGC was supported by the NIH/NIA grants: U01 AG032984, U24 AG021886, U01 AG016976, and the Alzheimer’s Association grant ADGC–10–196728.

## Supplementary Tables

Supplementary Table 1. Characteristics of the cohorts used in the analysis.

Supplementary Table 2. Summary of homozygosity measures for each individual study and the merged data set, considering two minimal ROH length cut-offs, 1 Mb and 1.5 Mb.

Supplementary Table 3. Summary statistics for the difference in homozygosity measures calculated using two different methods

Supplementary Table 4. Effect of genome-wide homozygosity measures in Alzheimer’s disease for each individual data set

Supplementary Table 5. Effect of genome-wide homozygosity measures in Alzheimer’s disease for the joint analysis, excluding deletions.

Supplementary Table 6. Effect of genome-wide homozygosity parameters in Alzheimer’s disease for the joint analysis, considering the effect of clonal mosaicism in aged populations.

Supplementary Table 7. Consensus ROHs associated with Alzheimer’s disease in the whole dataset.

Supplementary Table 8. Gene-based results for genes located in consensus ROHs associated with Alzheimer’s disease in the whole dataset.

Supplementary Table 9. Demographics for the pool of inbred individuals.

Supplementary Table 10. Consensus prioritized ROH based on the map of inbred Alzheimer’s disease patients.

Supplementary Table 11. Gene-based results for genes located in consensus prioritized ROH based on the map of inbred Alzheimer’s disease patients.

Supplementary Table 12. Variant annotation and functional effect prediction.

## Supplementary Figure Legend

Supplementary Figure 1. Quality control for A) ancestry and B) relatedness in the exome. All possible pairs had Pi-hat < 0.1875, a Z0 ≥ 0.75 and a Z1 ≤ 0.25.

Supplementary Figure 2. Boxplot for FROH per individual at ROH calling with 1Mb and 1.5Mb. Red line represents FROH = 0.0156 (mean inbreeding coefficient for kinship of second cousin marriage).

Supplementary Figure 3. Mean number of ROHs versus mean total sum of ROHs in Mb for the 10 cohorts explored, according to different ROH calling parameters. A) ROH length set to 1 Mb. ROH calling conducted with different number of markers per data set; B) ROH length set to 1.5 Mb. ROH calling conducted with different number of markers per dataset; C) ROH length set to 1 Mb. ROH calling conducted with the fraction of markers shared between data sets (2.6M); D) ROH length set to 1.5 Mb. ROH calling conducted with the fraction of markers shared between data sets (2.6M).

Supplementary Figure 4. Violin plots showing the distribution of ROH > 1.5 Mb within each data set and in the merged data for the homozygosity parameters (NROH, SROH, AVROH, FROH).

Supplementary Figure 5. Violin plots showing the distribution of ROH > 1.5 Mb within each data set and in the merged data for the homozygosity parameters (NROH, SROH, AVROH, FROH), split by case control status.

Supplementary Figure 6. Transformed distribution for the homozygosity measures. Transformation was performed using an inverse rank normal transformation with the “rankNorm” option in the RNOmni package in R.

Supplementary Figure 7. Distribution for average length of individual ROH segments for: non-inbred individuals (FROH < 0.0156), second-degree relatives (FROH > 0.0156) and first-degree relatives (FROH > 0.0625).

---

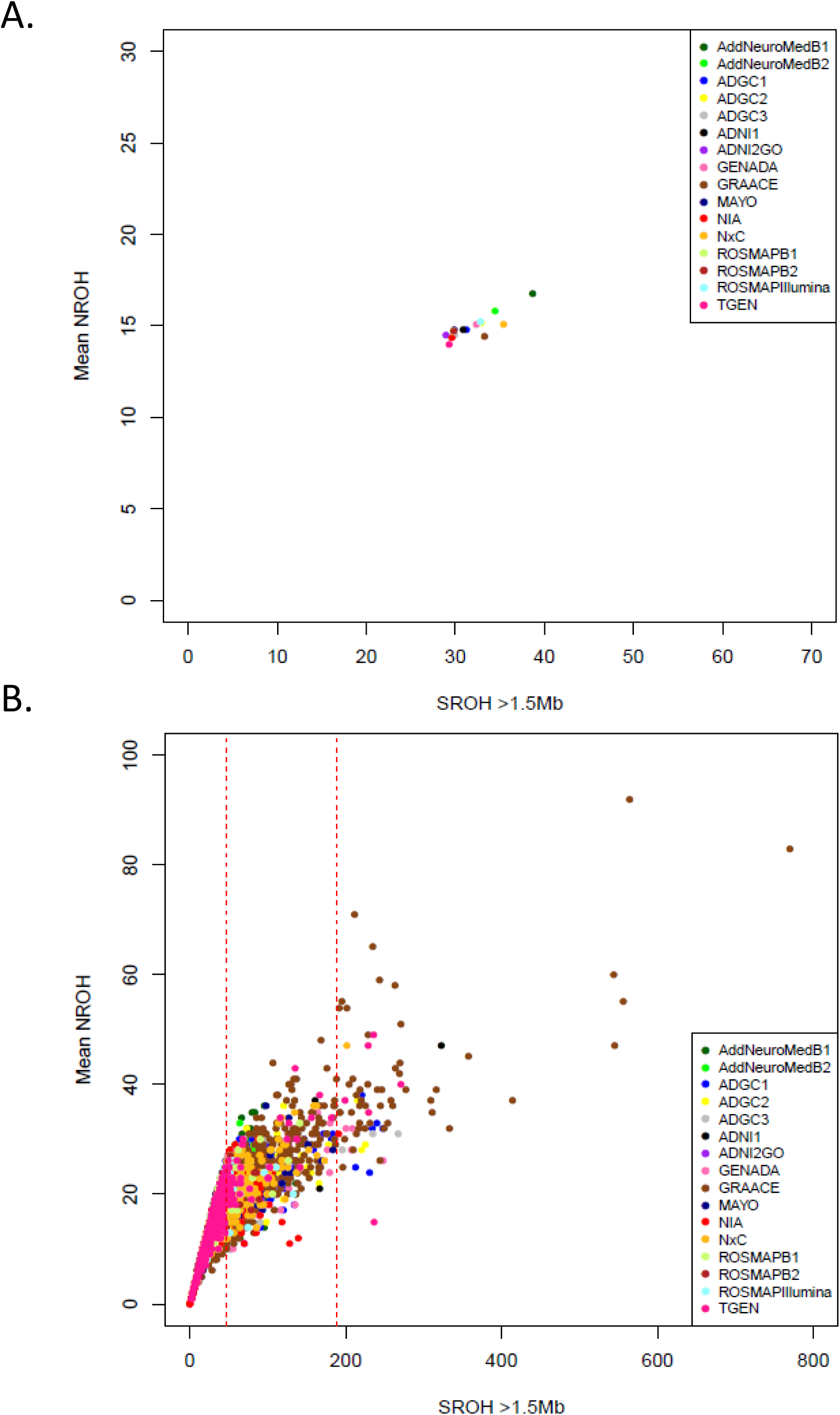

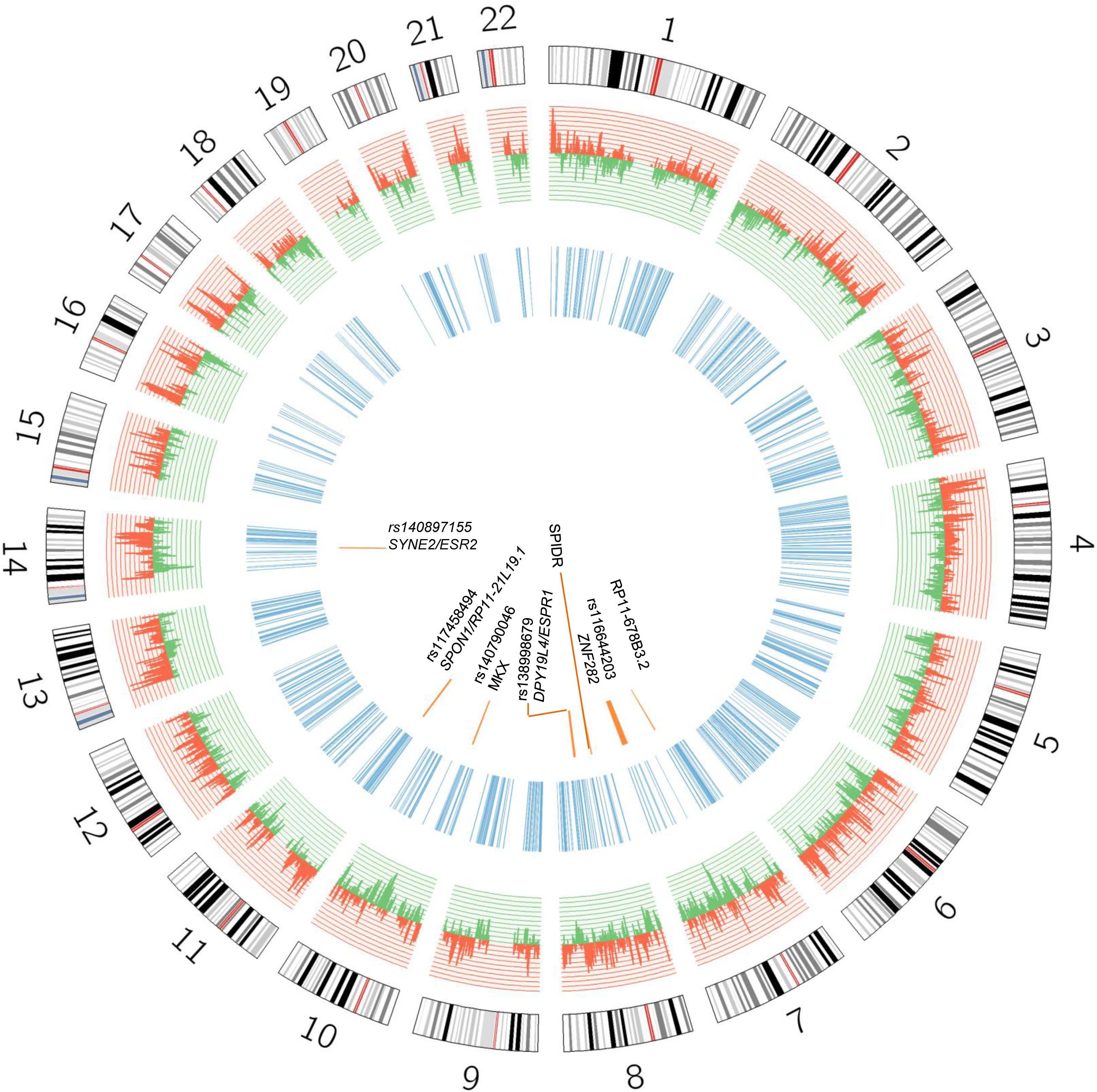

## Notes

Conflict of Interest: None.

