## Supplementary Figures for "Autosomal Recessive Alzheimer’s disease (arAD): homozygosity mapping of genomic regions containing arAD loci": Supplementary_Figure_7.pptx

### Slide 1
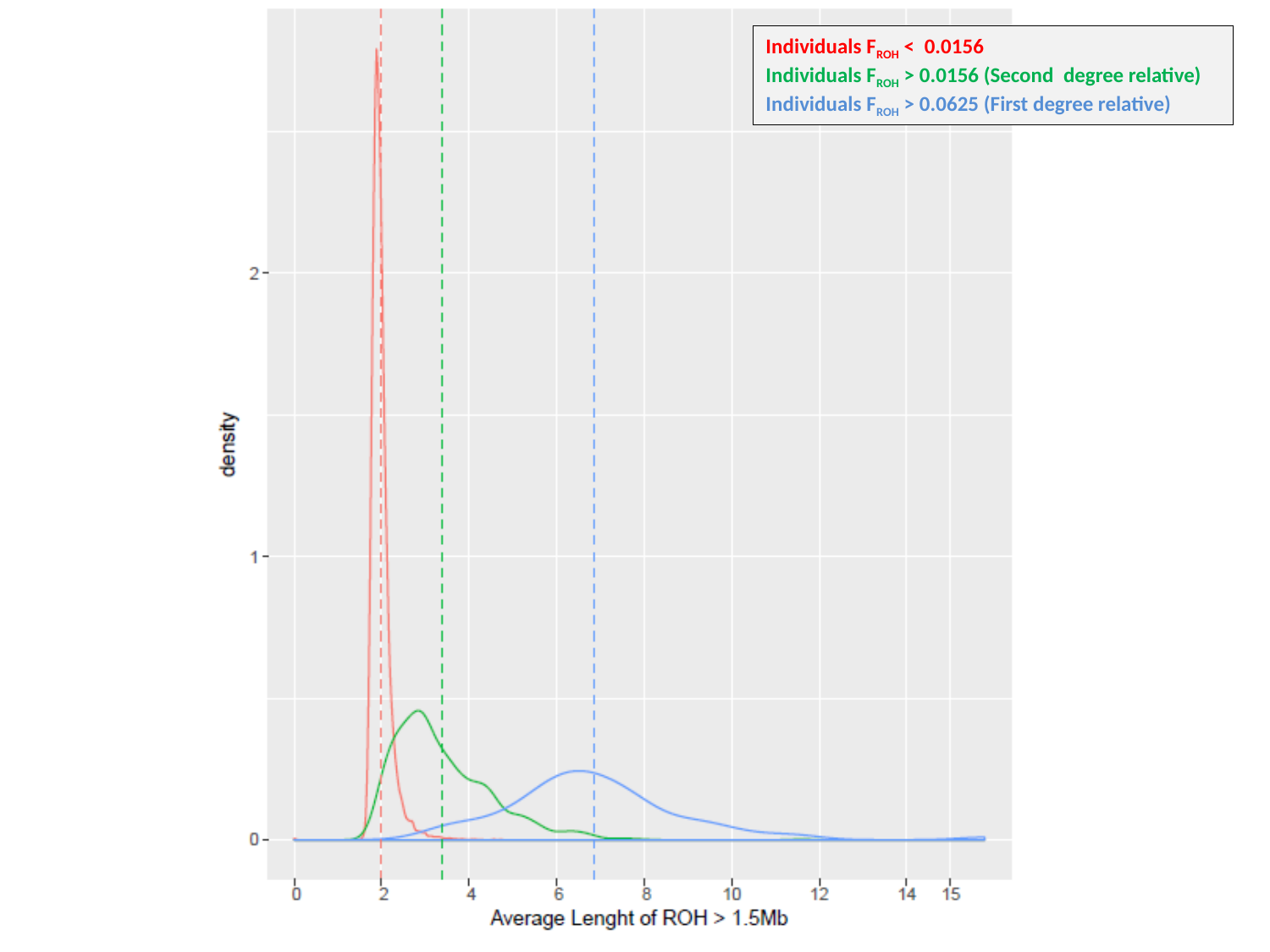

Individuals F_ROH_ < 0.0156
Individuals F_ROH_ > 0.0156 (Second degree relative)
Individuals F_ROH_ > 0.0625 (First degree relative)
