## Supplementary figures and images for "Autosomal Recessive Alzheimer’s disease (arAD): homozygosity mapping of genomic regions containing arAD loci"

### Supplementary_Figure_1.pptx

#### Slide 1
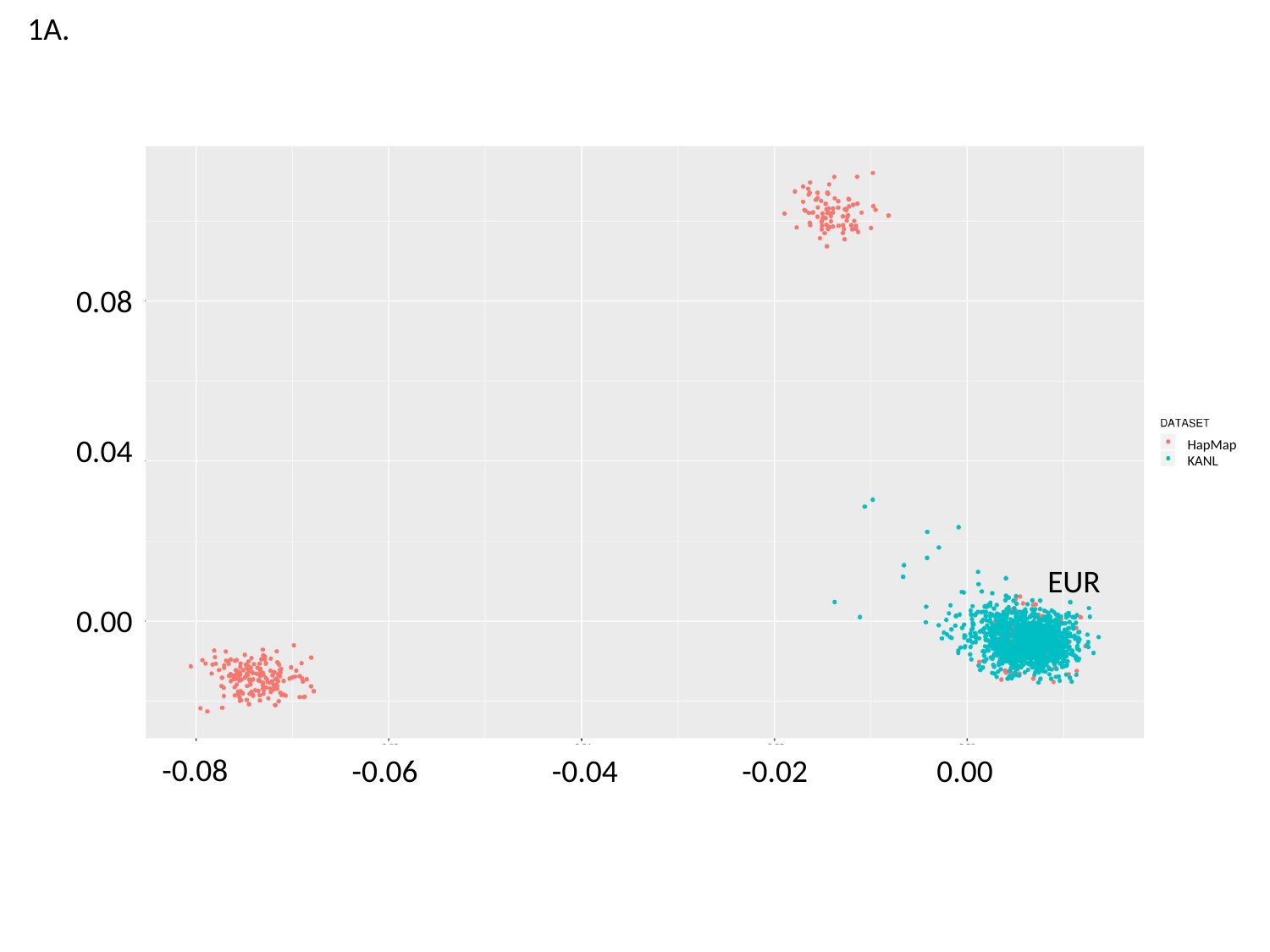

1A.
0.08
0.04
HapMap
KANL
EUR
0.00
-0.08
-0.06
-0.04
-0.02
0.00

#### Slide 2
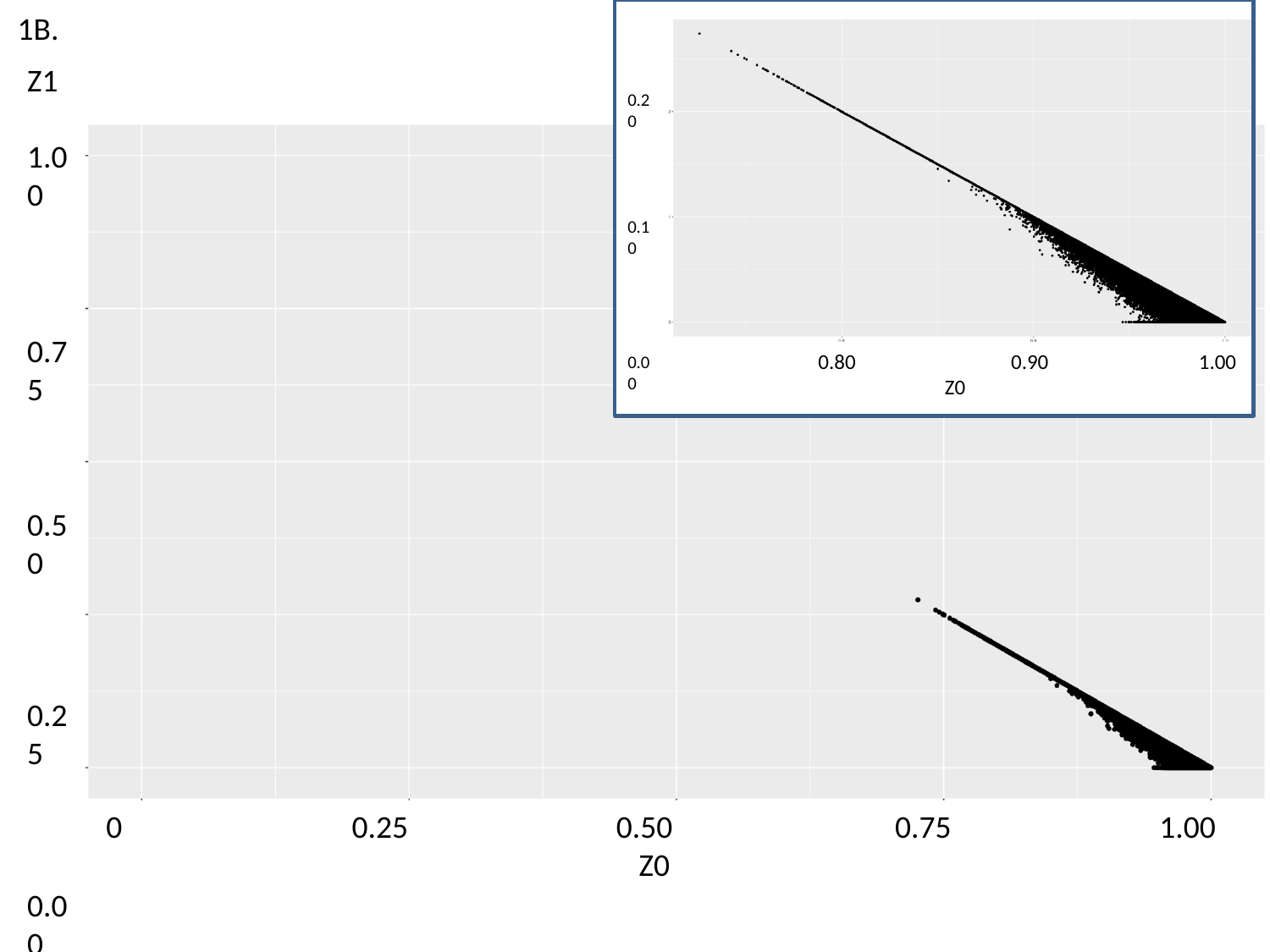

1B.
0.20
0.10
0.00
 0.80 0.90 1.00
 Z0
Z1
1.00
0.75
0.50
0.25
0.00
 0 0.25 0.50 0.75 1.00
 Z0

### Supplementary_Figure_2.pptx

#### Slide 1
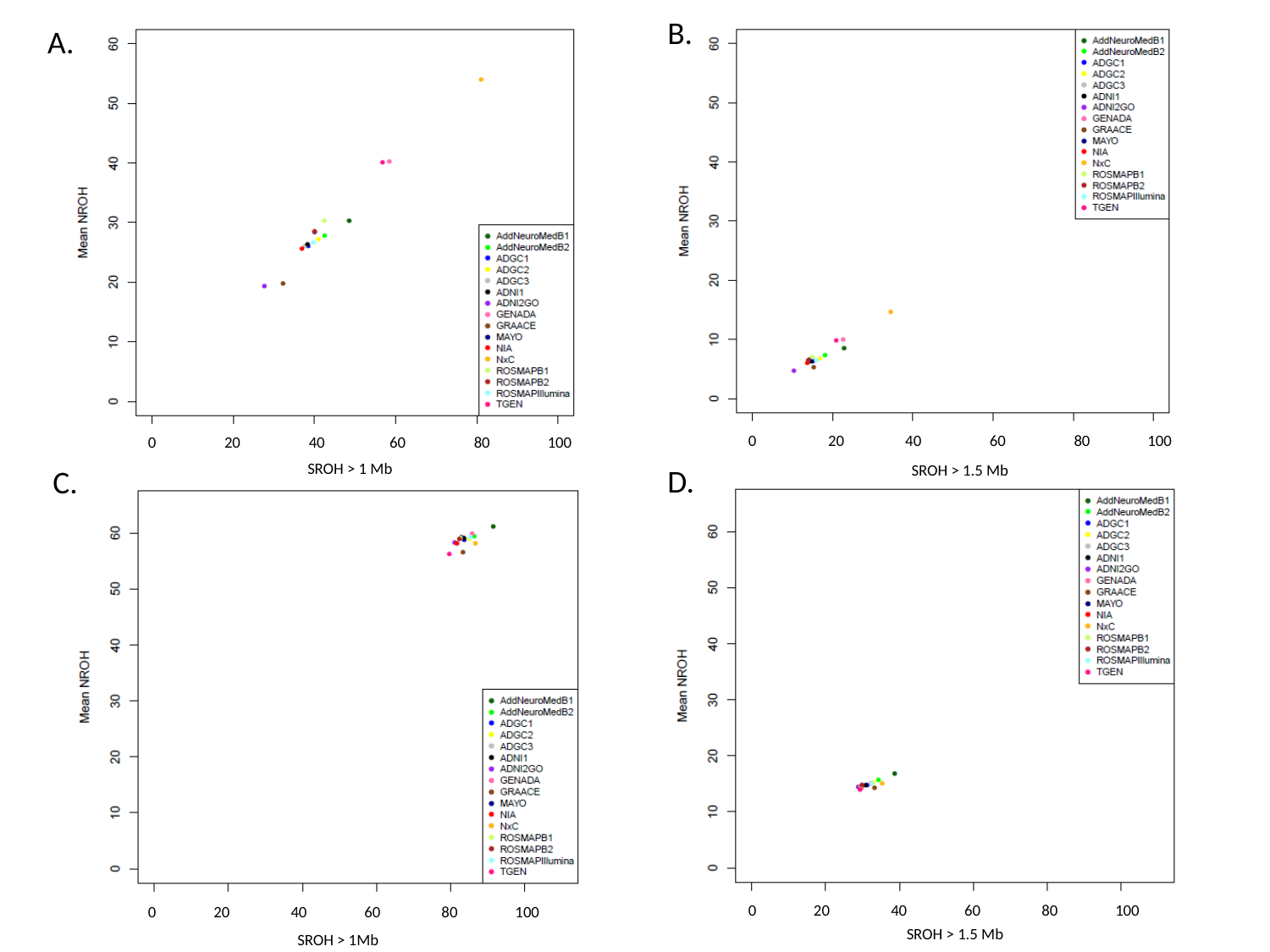

B.
A.
 0 20 40 60 80 100
 0 20 40 60 80 100
SROH > 1 Mb
SROH > 1.5 Mb
D.
C.
 0 20 40 60 80 100
 0 20 40 60 80 100
SROH > 1.5 Mb
SROH > 1Mb
SROH > 1Mb

### Supplementary_Figure_3.pptx

#### Slide 1
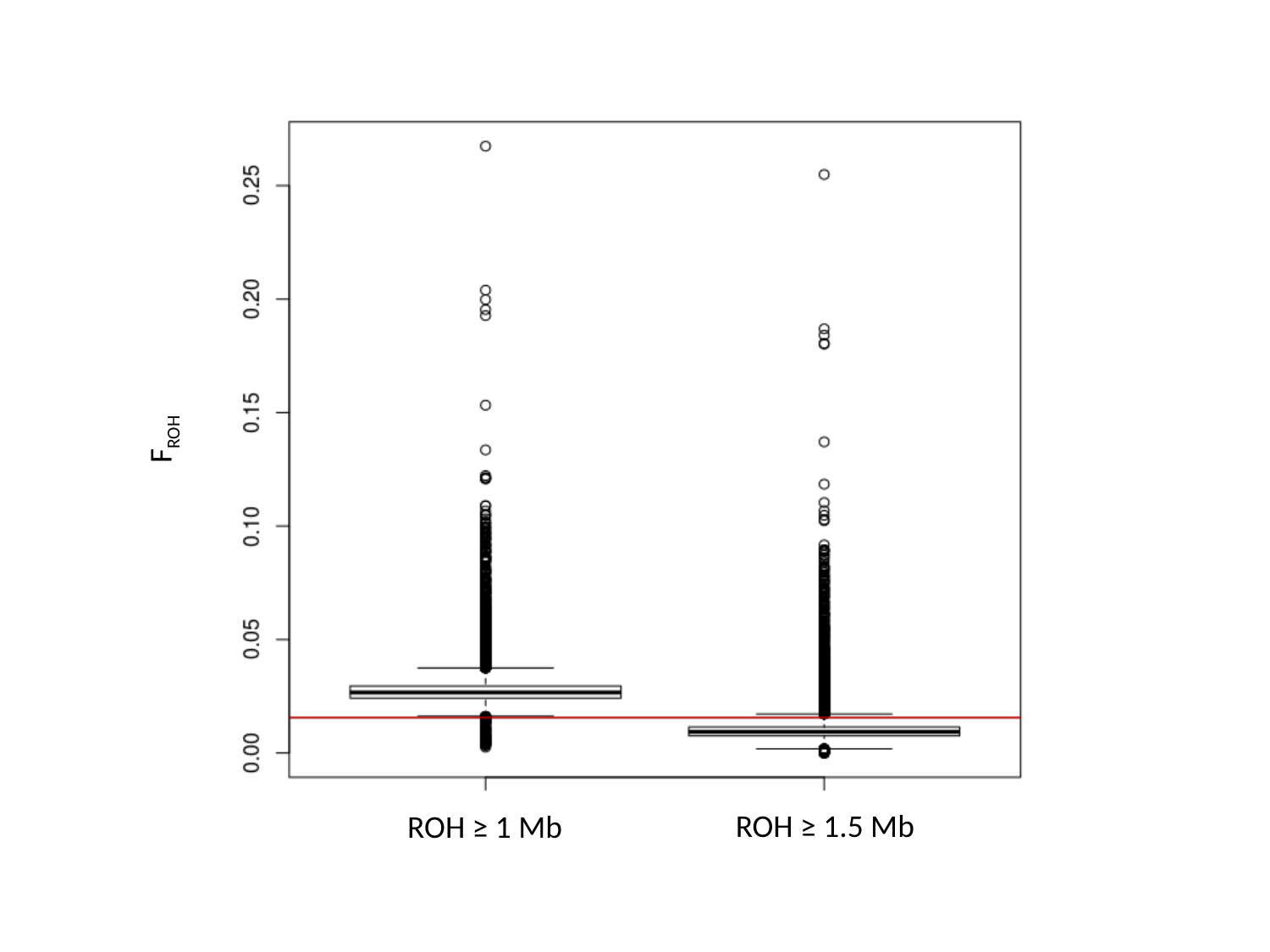

F_ROH_
ROH ≥ 1.5 Mb
ROH ≥ 1 Mb
ROH ≥ 1.5 Mb
ROH ≥ 1 Mb

### Supplementary_Figure_4.pptx

#### Slide 1
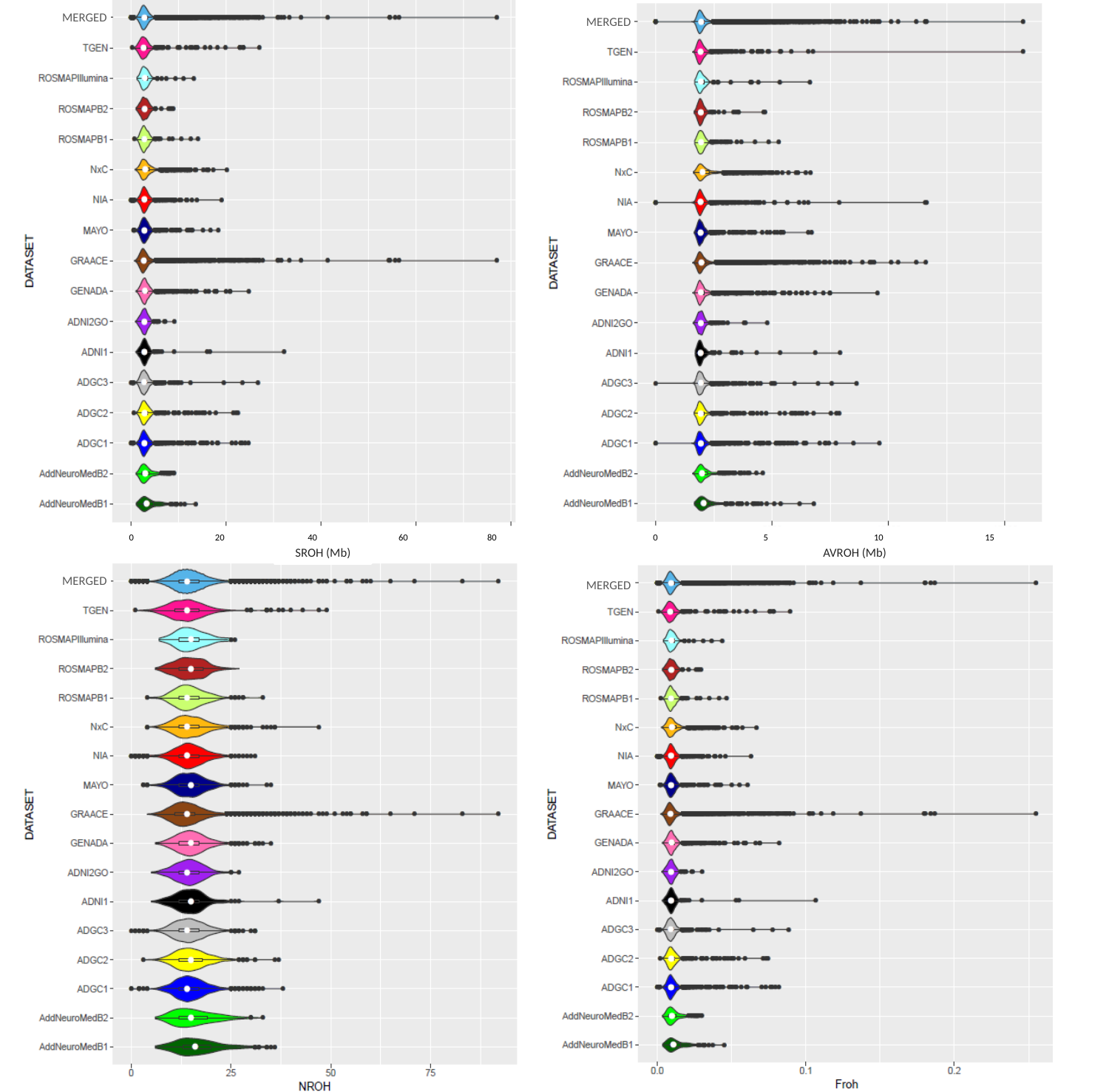

MERGED
MERGED
MERGED
 0 20 10 15
 0 20 40 60 80
 0 5 10 15
AVROH (Mb)
SROH (Mb)
AVROH (Mb)
MERGED
MERGED
MERGED
MERGED

### Supplementary_Figure_5.pptx

#### Slide 1
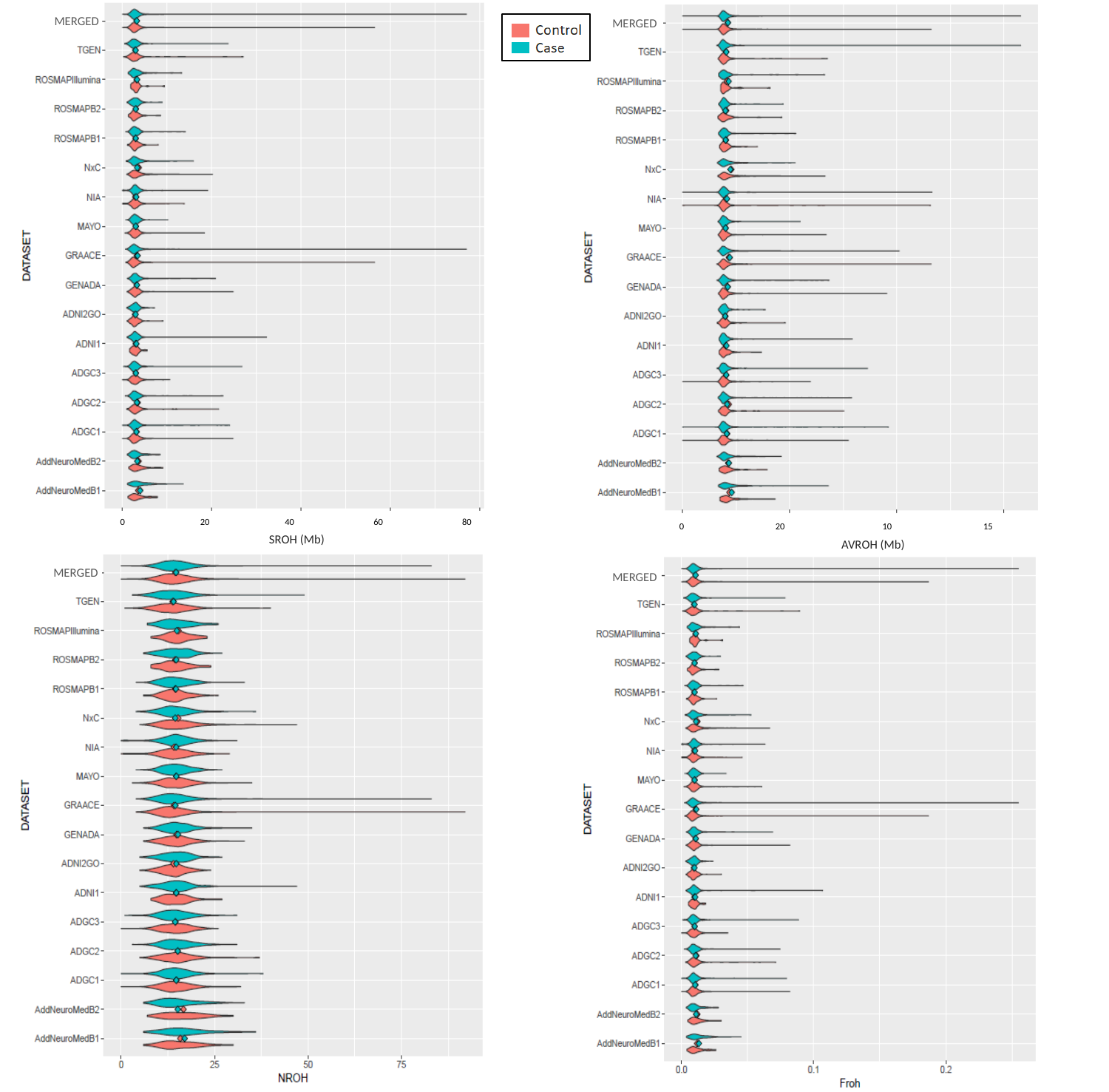

MERGED
MERGED
 0 20 40 60 80
 0 20 10 15
SROH (Mb)
AVROH (Mb)
MERGED
MERGED

### Supplementary_Figure_6.pptx

#### Slide 1
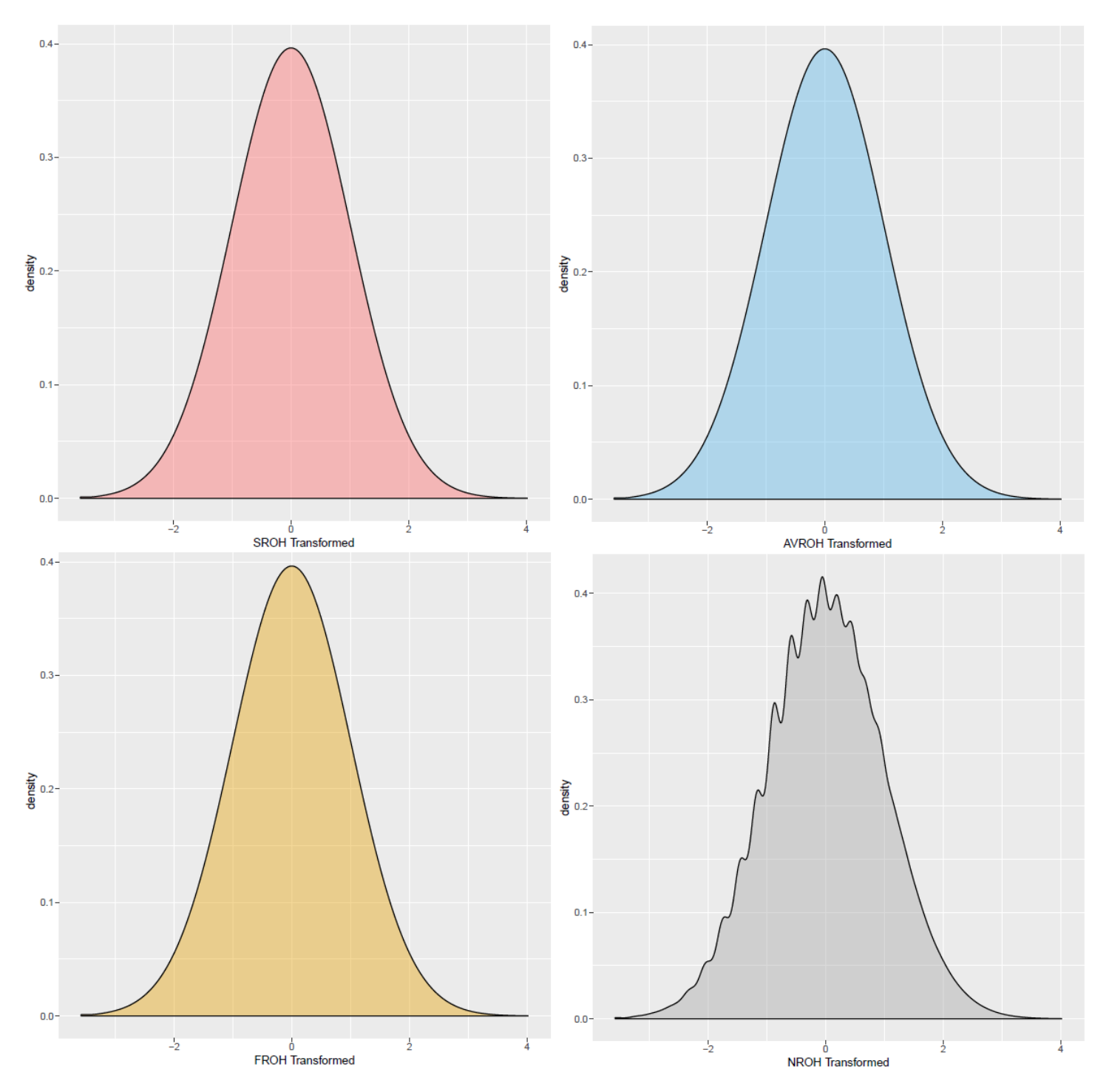
